# Unexpected D-tour Ahead: Why the D-Statistic, applied to Humans, Measures Mutation Rate Variation and Not Neanderthal Introgression

**DOI:** 10.1101/2024.12.31.630954

**Authors:** William Amos, Eran Elhaik

## Abstract

The D statistic is widely used to infer introgression and played a pivotal role in establishing as fact that humans interbred with Neanderthals, leaving a lasting genetic legacy. However, D relies on two untested assumptions: mutation rate constancy between groups and a negligible contribution from recurrent mutations. Together, these preclude a previously ignored alternative explanation of the data based on mutation-rate variation across human populations. Here we show that neither assumption is valid and infer that essentially all sites contributing to non-zero D carry recurrent mutations. When D is calculated separately for different sequence motifs, it varies far more than can be explained using a model based on introgressed haplotypes, where non-independence due to being located on the same haplotype largely prevents different motifs from generating different values of D. Large variation in D between triplets appears only possible if D is driven by processes acting at the level of individual bases. We further show that D is dominated by sites where the Neanderthal allele matches the human major allele and, more specifically, where the Neanderthal allele is fixed in humans outside Africa. This is the exact opposite of expectations under the introgression hypothesis, where Neanderthal alleles should mostly be rare and found only outside Africa. Finally, we tested a new model in which heterozygosity modulates mutation rate. Remarkably, heterozygosity differences, relative mutation rate, and D are all strongly inter-correlated, not only in African – non-African comparisons but also in comparisons among African and among non-African population pairs, and even across the genome. We conclude that the mutation rate variation hypothesis provides a more compelling explanation for the observed patterns in human-Neanderthal genetic relationships than the introgression hypothesis and that the concept of archaic introgression into humans and its implications for hominin-derived traits must be rethought.

## Introduction

The idea that humans outside Africa carry a small but ubiquitous legacy that results from interbreeding with Neanderthals sometime around 40 – 50,000 years ago was first proposed 15 years ago when the draft sequence of the Neanderthal genome was published (Green et al. 2010) and quickly became accepted as fact (Prüfer et al. 2014; Sankararaman et al. 2014; Skov et al. 2020). Once established, the introgression narrative led to wide ranging searches for links to human traits, and many papers appeared reporting how Neanderthal or Denisovan sequences contributed to, for example, adaptation to high altitude living in Tibetans (e.g., Huerta-Sánchez et al. 2014); anthropometric traits such as lip thickness (Bonfante et al. 2021) and sitting height (Dannemann and Kelso 2017); and immune-related phenotypes such as genetic risk factors for severe COVID-19 (Zeberg and Pääbo 2020).

There is no dispute that non-Africans are genetically closer to Neanderthals compared with Africans, regardless of how this is quantified (Green et al. 2010; Skov et al. 2020). However, this pattern can arise in two different ways (Fig. 1A). One explanation is that Neanderthals and non-Africans are unexpectedly *similar* because non-Africans carry segments of Neanderthal DNA. An alternative possibility is that Neanderthals and Africans are unexpectedly *different* because Africans have a relatively higher mutation rate (Amos 2020; Amos 2021). We refer to the latter as the mutation rate variation hypothesis (MRVH). The vast literature on archaic introgression into humans all follows the original study in assuming, without testing, that mutation rate is constant across human populations (e.g., Patterson et al. 2012; Sankararaman et al. 2014; Lopez Fang et al. 2024). This critical assumption effectively bypasses the MRVH and forces all downstream analyses to conclude that introgression has occurred.

**Figure 1.**
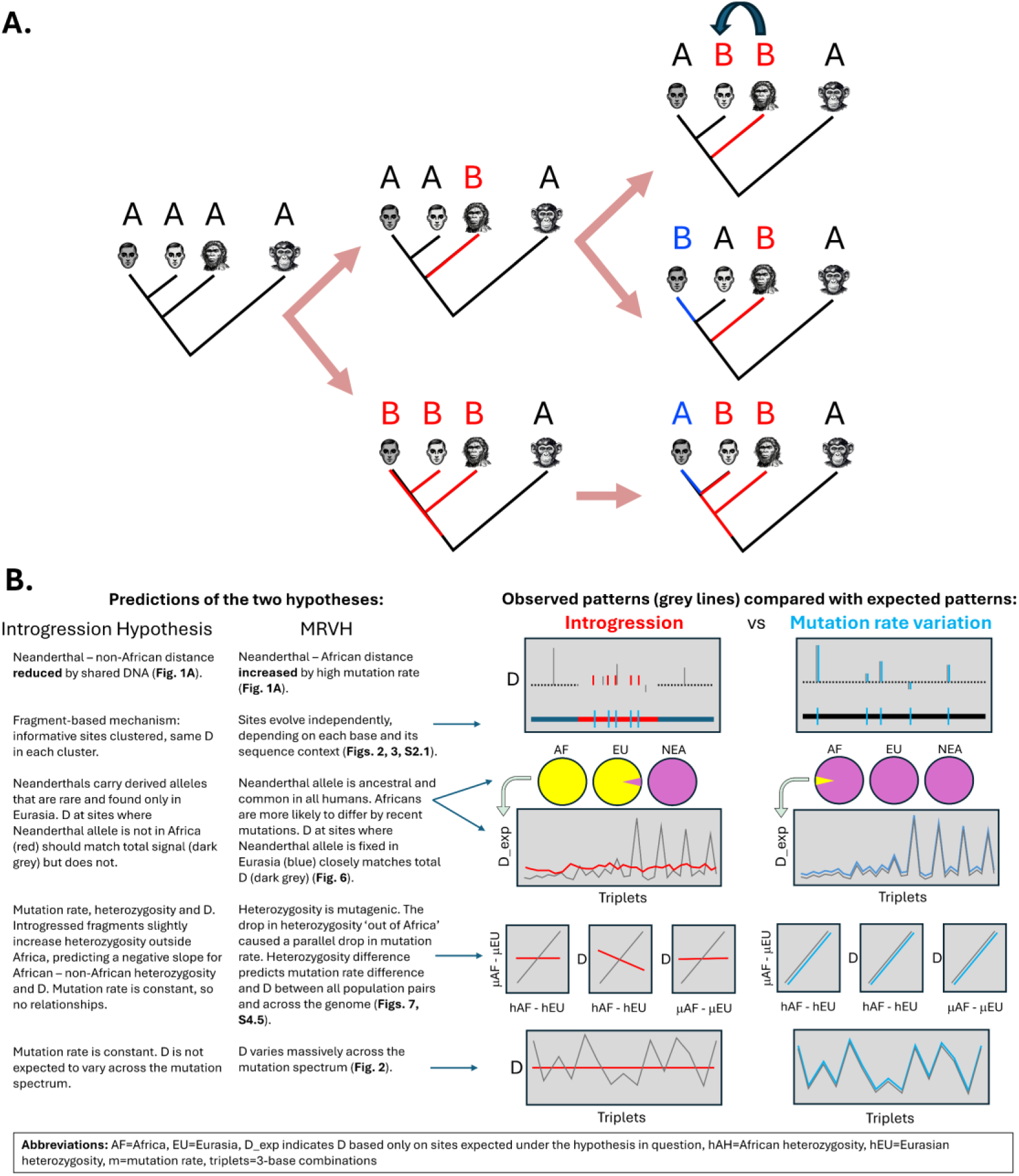
Comparison of two hypotheses explaining the apparent genetic proximity of Neanderthals to Europeans relative to Africans and their predictions. A) Illustrated is a single site where the ancestral state carried by all taxa is ‘A’. Under introgression (top row), an A->B mutation occurs in Neanderthals and enters Europeans via interbreeding to form an ‘ABBA’ pattern. Under mutation rate variation, both ABBAs and BABAs can be created by recurrent mutations (blue), but a higher mutation rate in Africa means that the excess pattern is generated by an African mutation. In the middle row, A->B mutations in Africa create BABAs, while in the bottom row, B->A mutations create ABBAs. In this way, the same mutation rate difference creates opposing signals that depend on which taxa carry the first mutation (red). Recurrent mutations in Europeans also create opposing signals at a lower rate. Branch lengths are not drawn to scale. B) A comparison of the major predictions of two hypotheses tested in our study (left side) and the agreement of those predictions with the data (right side). The data are fully consistent with the predictions of MRVH, rather than predictions made by introgression.

The simplifying assumption of mutation rate constancy is so widely used in population genetic theory that it is easy to forget it is a null assumption with no empirical basis. Direct estimates of mutation rate remain challenging, even with high-coverage genome sequences from many families (Kong et al. 2012b; Jónsson et al. 2017; Porubsky et al. 2025), so comparative rate estimates rely on indirect methods that carry their own assumptions. However, when mutation rate variation has been looked for, it has often been found (Kong et al. 2012a) varying among primate species (Brown et al. 1982; Chintalapati and Moorjani 2020) between human populations (Amos 2013; Mallick et al. 2016; Harris and Pritchard 2017; Garcia-Salinas et al. 2025) and, massively, along a chromosome (Rogozin and Pavlov 2003; Nesta, Tafur, and Beck 2021). Interestingly, in the few cases where mutation-rate variation among populations is considered alongside introgression as competing explanations for why non-Africans are closer to Neanderthals, the data appear to fit MRVH better than introgression (Amos 2020; Amos 2021). The mutation-rate-constancy assumption has a complementary counterpart: the assumption that recurrent mutations occur at a negligible rate, so shared variants can safely be interpreted as having a common origin.

Among many diverse approaches that attempt to quantify interbreeding (see review by Gopalan et al. 2021), one of the most widely used is based on a simple statistic, D, that supposedly captures the extent to which one human or human population is genetically closer to Neanderthals (or Denisovans) than to another human (Durand et al. 2011; Rogers and Bohlender 2014; Peter 2016). Along with most other methods (e.g., Racimo et al. 2015; Schaefer, Shapiro, and Green 2021), D is founded on the twin assumptions discussed above, that mutation rate does not vary among human populations and that recurrent mutations are sufficiently rare that they can reasonably be ignored following the infinite-sites model of mutation (Green et al. 2010; Durand et al. 2011; Sankararaman et al. 2016). Under these assumptions, there are only two main ways in which a polymorphism can exist both within humans and outside humans: introgression and incomplete lineage sorting (ILS) (Blom, Hipsley, and Zhang 2026). ILS occurs when the genealogical history of a particular stretch of DNA is older than the split between the species being studied. Here, ILS refers to a state where the root of the human gene tree predates the human-Neanderthal split, allowing one or more human lineages to root within Neanderthals rather than humans.

As we noted, the first assumption of mutation-rate constancy across populations has no robust empirical basis. Indeed, available evidence suggests that, although mutation rates are difficult to measure, they do vary. The second fundamental assumption, that recurrent mutations are vanishingly rare, is also questionable. As evidenced by dbSNP data (Sherry et al. 2001), human single nucleotide polymorphisms (SNPs) often carry three or even all four bases. Unless due to sequencing errors, such multiallelic sites must have experienced at least two mutations since the most recent common ancestral base. Even after allowing for the huge sample sizes in dbSNP, where extremely rare alleles are seen that would have been missed in smaller data sets, recurrent mutations appear far commoner than many assume. Because both assumptions underpinning the introgression statistic appear unsound, we looked more carefully at what drives D. In particular, we focus on differences in the predictions of the two competing hypotheses (Fig. 1B, left panel). Through this, we aim to better understand what exactly D captures in humans and the likely downstream implications for our understanding of archaic Hominin introgression. We use published, high-coverage genome sequences alongside extensive simulations to run a series of tests designed to expose any possible influence of mutation-rate variation on allele-sharing rates between the taxa of interest. We find that, in all cases, the MRVH fits the data better, often leaving little room for an appreciable signal due to introgression.

## Results

To better understand the signal driving D, we analysed the high-coverage 1000 genomes (1KG) (2504 unrelated individuals drawn from 26 global population groups) (Clarke et al. 2016) aligned to three high-coverage archaic genomes, the Altai (Prüfer et al. 2014) and Vindija (Prüfer et al. 2017) Neanderthals and the Denisovan (Peyrégne et al. 2025). We begin by revisiting the two key assumptions on which D is founded. We then adopt two strategies to determine whether positive D truly measures introgression. First, we exploit the fact that different sequence motifs have widely varying inherent mutation rates. This allows us to compare predictions from the introgression hypothesis, where Neanderthal variants should contribute to D in proportion to their genome-wide frequencies, with predictions from the MRVH, where different motifs may contribute very differently depending on their relative mutability. Second, we partition D by the characteristics of the contributing sites. Critically, we compare D calculated from sites where the Neanderthal allele is rare in humans, which must include the overwhelming majority of sites generated by introgression, with D calculated using only sites where the Neanderthal allele is fixed in non-Africans, these being highly unlikely to be the product of introgression. Throughout our study, we compare our empirical results with outputs from realistic forward simulations in which the mutation rate is either held constant or varies over time and/or across populations. For realistic demographic parameters, we use those from the SLiM manual (Haller, Ralph, and Messer 2026) based on work by Gravel et al. (2011) (“the Gravel model”).

We use the following terminology. ‘Human’ always refers to Homo sapiens and not to related species of Homo. The ‘standard model of introgression’ was taken as a single main interbreeding event around 50,000 years ago, after some humans left Africa but before they colonised the rest of the world (Green et al. 2010; Durand et al. 2011). Sites counted as ABBA or BABA states are called ‘D-informative.’ D-informative sites comprise two classes of site: (i) ABBAs and BABAs generated in equal numbers through incomplete lineage sorting (ILS); and (ii) an excess of either ABBAs or BABAs that drives non-zero D. We refer to these as D-symmetric and D-asymmetric sites, respectively, though it is rarely possible to assign any individual site to one or the other. However, these categories do yield different expectations for the aggregate distribution of ABBA and BABA counts under competing evolutionary models. We refer to the Vindija and Altai Neanderthals as Vindija and Altai. Unless specified otherwise, ‘D’ refers to D(Yoruba, French, Altai, chimpanzee). We extend the ABBA-BABA nomenclature by using ‘P’ to indicate a base that is polymorphic in humans, and we use a constant order: human / Vindija / Altai / Denisovan / chimpanzee. Thus, PAAAA indicates a site that is polymorphic only in humans. We use ‘constant mutation rate’ specifically to refer to scenarios where mutation rate for any given class of variant does not vary between different human populations. In several analyses we explore the mutation spectrum, *sensu* Harris and Pritchard (2017) and Harris (2015), where mutations occur at different rates for the central base in three-base combinations (triplets), most notably where the central base is the C of a CG pair (XCG triplets).

### Challenging the assumptions used to infer introgression

#### Recurrent mutations are not vanishingly rare and must not be ignored

We began by counting SNPs with three alleles and comparing their frequency to the number of sites that drive D. The numbers of both classes of site are strongly dependent on sample size. To avoid possible sequencing errors, we counted only sites where the rarest allele was present in at least three copies. Sites carrying three genuinely different bases must reflect two independent mutations occurring since the most recent common ancestral base: to generate three different alleles, at least one of the mutations must have been a transversion.

Across the autosomal genome, in the 1KG data we counted 611,742 sites that carry three bases. Since transition (TS) mutations typically occur at twice the rate of transversions (TV), for each site with three alleles we can expect twice this number of similar but ‘silent’ sites where two transitions create two, rather than three alleles (Segurel, Wyman, and Przeworski 2014). This gives us around 1,800,000 sites carrying recurrent mutations in a 3Gb genome or one every 1.67kb. For D, we count around 300,000 informative sites, depending on the population combination used, of which 16,400 are D-asymmetric (ABBA minus BABA). This equates to roughly one D-asymmetric site every 190kb. Thus, far from occurring at a negligible rate, we find there are ∼100 sites that carry a recurrent mutation for each D-asymmetric site.

In the base simulation, 0.9% of sites that are polymorphic in a sample of 50 ‘African’ and 50 ‘non-African’ humans carry three or more alleles, and 1.9% carry ‘silent’ recurrent mutations (i.e., only two alleles but more than one mutation event). This gives a recurrent visibility ratio of 2.06:1, close to the expected ratio of two, calculated assuming transversions occur at twice the rate of transitions. In terms of density along a chromosome, our simulations generate a human genome equivalent of 919,560 sites with three or more alleles, similar to the observed rate, plus 1,883,310 ‘silent’ sites. Sites carrying recurrent mutations occur at an average density of 0.85 sites per kilobase (approximately one every 1177 base pairs). Our simulations suggest that an individual haploid human genome will carry 1113 variants created independently by recurrent mutations in Neanderthals and 18,642 variants that are recurrent with respect to the chimpanzee. The likelihood of observing recurrent mutations increases with the number of chromosomes sampled. Our simulations predict that a sample of 100 African and 100 non-African chromosomes would include 9,220 sites carrying at least one recurrent Neanderthal variant and 172,920 sites with at least one recurrent chimpanzee variant.

#### Recurrent mutations dominate the signal captured by D in humans

The number of mutations at each informative site fundamentally distinguishes the two competing hypotheses. Under introgression, D informative sites generally carry a single mutation: recurrent mutations are not integral to the mechanism driving D and are included only by chance. By contrast, recurrent mutations play a central role in the MRVH and should be carried by every D-asymmetric site. One way to assess the frequency of recurrent mutations is through TS:TV ratios. Across the whole genome, the TS:TV ratio is widely reported to be approximately two (The International HapMap Consortium 2003; DePristo et al. 2011), so the mutation rate of transitions, mTS, is twice the mutation rate of transversions, mTV, i.e., m_TS_ = 2m_TV._ However, the probability of a recurrent mutation scales with mutation rate squared, giving equivalent probabilities for recurrent TS and TV mutations of m_TS_^2^ and 4m_TV_^2^, respectively. Thus, the TS:TV ratio should be around two at sites carrying single mutations but will increase to about four at sites with recurrent mutations. Analysing the 1KG data, we show that the ratios are indeed ∼4 at sites that have most likely experienced recurrent mutations, specifically sites where all non-human hominins carry the human reference allele and the chimpanzee carries the alternate allele when this is also the human minor allele ([aR]RRRA, where R = reference, A = alternate and lower cases are the human minor allele, TS:TV ratio = 3.98). Moreover, partitioning the data by mutating triplet reveals how the average ratio of 3.98 embraces huge variation, from below 1.5 (e.g. AAA, AAC) up to above 10 for triplets of form [any base]CG (= ‘XCG’ triplets), with a maximum ratio of 22.3 for ACG (ESM1, Table S1.1).

Having confirmed that sites most likely to carry recurrent mutations have close to the expected TS:TV ratio of four, we next considered the TS:TV ratios at sites that either do or do not contribute to D. Sites contributing to D have an overall ratio of 2.79 compared with 2.23 for sites that do not. Interpreting 2.79 as a mixture of sites with ratios of 2.23 and 3.98 suggests that 32% of D-informative sites carry recurrent mutations. This calculation is very approximate because it is hampered both by observation biases and because recurrent mutations will occur at different and unknown rates in all classes of site. For example, the overall ratio for [rA]AAAR might be considered equivalent to [aR]RRRA just with R and A reversed, but the ratio is much lower, at 2.67. We believe this reflects an observation bias, as the reference allele is unlikely to be extremely common. Consequently, [rA]AAAR sites will tend to be enriched for sites where R and A are at similar frequencies and these are much more likely to be the product of a single mutation followed by ILS, hence the lower TS:TV ratio.

#### D varies strongly across the mutation spectrum, consistent with mutation rate variation among human populations

The next key difference between introgression and the MRVH is the assumption of mutation rate constancy. Durand et al. (2011) show analytically that if mutation rate does not vary between populations, D will not vary, even if mutation rates vary between sites: higher mutation rates generate more derived variants in Neanderthals that then become introgressed alleles if interbreeding occurs, but the higher rate also creates more sites that generate shared polymorphism through ILS. However, Durand et al. modeled mutation rate as a first-order term and ignored recurrent mutation and the probability that it increases with the square of the mutation rate, so high-mutation-rate sites contribute disproportionately to ABBA–BABA asymmetry. When recurrent events are ignored, mutations make no detectable net contribution to *D*, an assumption that breaks down under the MRVH.

In empirical data, D varies dramatically between triplets, ranging from around zero up to approaching 0.12 in typical African – non-African population comparisons (Fig. 2). In general, the peaks and troughs of every profile tend to occur at the same triplets such that most of the profile variation involves shifts on the Y-axis, with small shifts between population comparisons within a regional comparison (lines of the same colour) and larger shifts between different regional comparisons (different colours). The most striking differences are between comparisons that do or do not involve Africans. A model based on introgression can only drive these patterns if selection modulates the fate of haplotypes based on the variants they carry, but this is biologically impossible because selection acts on base substitutions, not on the triplet context: the 64 classes of triplets we present cover just four different types of substitution: transitions and transversions at A and C.

**Figure 2.**
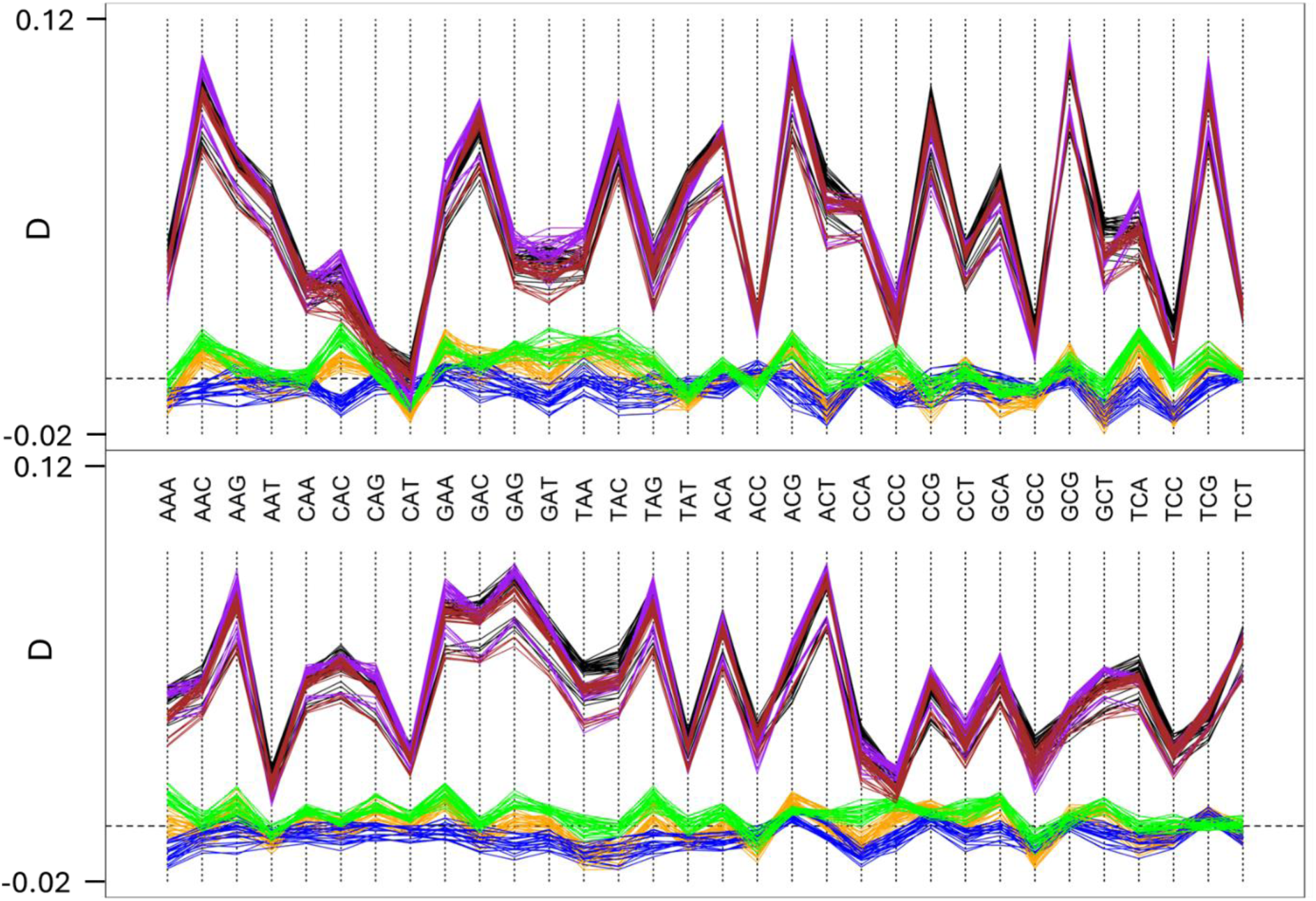
Variation in D by triplet. Triplets were defined as the ancestral state, assuming the human minor allele is the derived state. D is calculated for each triplet separately, partitioned into transversions (top panel) and transitions (bottom panel). All possible pairwise comparisons are made between African (AFR), European (EUR), East Asian (EAS) and South Asian (SAS) populations: black (EUR-AFR); brown (SAS-AFR); purple (EAS-AFR); green (SAS-EAS); orange (EUR-EAS); blue (EUR-SAS). Every line is a different population comparison. D on the y-axis is the same in both panels and ranges from - 0.02 to 0.12.

Several other observations also rule out selection-based explanations. First, in purely non-African comparisons, D fluctuates above and below zero, a scenario that would require selection to consistently favour certain triplets in some regions and other triplets elsewhere. Second, while the profiles are generally parallel, for CAC transversions and a few other triplets, some profiles show peaks that correspond to troughs in others. This pattern would require selection to reverse direction over short timescales. Third, these patterns become even more extreme in CpG islands, known to exhibit distinctive context-dependent mutation rates (Carlson et al. 2018), with D rising to a massive 0.7 and dipping well below zero; these negative values occur even in African–non-African comparisons. Remarkably, the extreme CpG island profiles also differ greatly between chromosomes. Selection again does not offer a plausible explanation, since it would have to act far more intensely on introgressed material from CpG islands, and selection would also have to vary between chromosomes (for data and more discussion, see ESM2, Figs. S2.1, S2.2).

Taken at face value, and with selection excluded, variation in D across triplets would appear to prove that mutation rates vary between human populations: according to theory, if mutation rate is constant, D should not vary. However, our forward simulations reveal hidden complexities. On the one hand, the simulations confirm that, in the absence of mutation rate variation between populations and introgression, D does not differ from zero when calculated either over all sites (D=0.001) or for transversions (D = -0.003) or transitions (D = 0.002) separately (see ESM7, Table S7.1). On the other hand, when D is calculated for individual triplets, it *does* vary (Fig. S7.1), and the resulting values are positively correlated with equivalent values from empirical data (Fig. S7.2, S7.3). Although we do not yet fully understand the mechanism, the positive correlation between simulated and empirical values strongly suggests it is an emergent property reflecting a dynamic balance between triplet-specific mutation rates and the frequencies of different triplets.

Importantly, introducing a simulated mutation-rate drop between Africans and non-Africans increases the strength of the correlation between the empirical and simulated values and generates non-zero D (Fig. S7.3, Table S7.1). Across a range of simulated scenarios, a 50% drop in rate can give D up to 0.05, similar to empirical values typically in the range of 0.05–0.06. However, D depends strongly on the mutation model used. When the only variation in rate modelled is a simple TS:TV ratio of two, D is much lower than when the full inter-triplet variation is modelled. This is important because it emphasizes the key role of recurrent mutations, which are disproportionately likely at the most mutable sites: for any given average mutation rate, as variation in mutation rates across sites increases, so does the number of recurrent mutations. Moreover, although we have not modelled mutation hotspots and coldspots, these will further increase D (Aggarwala and Voight 2016; Koppetsch, Malinsky, and Matschiner 2024). We also modelled the impact of a general rise in human mutation rate, increasing the proportion of variants originating after the out-of-Africa event, which further increases D for any given drop in rate. In sharp contrast, the correlation between simulated and empirical triplet-specific D values disappears completely when introgression is added (Fig. S7.4). As before, data fit the MRVH unambiguously better than the introgression hypothesis.

#### D is driven by processes that act on individual bases, not fragments, precluding introgression as the dominant driver of variation in D

Under the introgression hypothesis, informative sites on the same introgressed fragment must contribute equally to D. This contrasts with the MRVH, where the mutation rates of individual bases are modulated by factors such as sequence context (Harris and Pritchard 2017; Koppetsch, Malinsky, and Matschiner 2024) and heterozygosity (Amos 2013), allowing bases to give high and low D despite being very tightly linked. This distinction is critical in light of the high degree of variability in D seen between triplets, because a mechanism based on fragments can only work if: (a) triplets are distributed highly non-randomly, with triplets that give similar D clustering together; and (b) selection either favours or purges introgressed fragments according to the triplets they carry. Neither of these conditions seems plausible. Nonetheless, we tested how non-randomly triplets are distributed. For this, we identified every instance of a given triplet that contributes to D and calculated expected D based on all D-informative triplets within 20kb (Fig. 2). All triplets give median Ds that are essentially the genome-wide average D of ∼5%. Crucially, D computed for that triplet alone often lies entirely outside the range of variation seen across all possible 40kb windows in the genome (Fig. 3). Thus, as expected, triplets are approximately randomly distributed with respect to each other. The widely differing D values seen for different triplets cannot be explained by a mechanism operating through fragments. We also tested whether a major source of heterogeneity within the genome, CpG islands, might play a role, finding that triplet-specific variation in D is exaggerated within islands (Figs. S2.1 and S2.2).

**Figure 2a.**
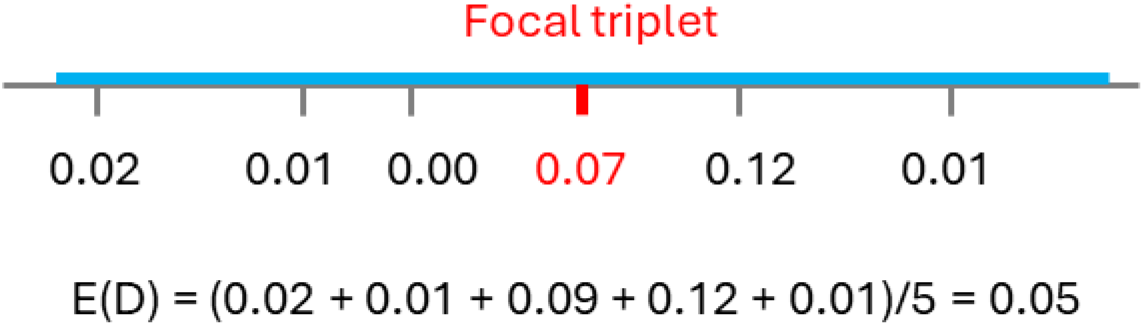
Calculation of expected D, E(D) for genomic fragments. All triplets contributing to D were identified. Each triplet was then treated as the focal triplet (red) and used to define the centre of a symmetrical 40kb window (±20 kb, in blue). We assigned each neighboring D-informative triplet within the window its genome-wide mean D value (black numbers), then averaged these values to estimate the expected D for the focal site, E(D). This procedure was repeated for every occurrence of every triplet type, generating a distribution of expected D values for each triplet. Under the introgression hypothesis, neighboring sites on the same introgressed fragment should share a common history and therefore exhibit similar D values, i.e., E(D) should closely match the observed genome-wide D of the focal triplet and the only way for D to vary between triplets would be for triplets to be strongly clustered according to the genome-wide D they give. By contrast, if D is determined primarily by the intrinsic mutational properties of individual triplets rather than by the surrounding genomic fragment, neighboring triplets will not predict the focal triplet’s D, and E(D) will remain close to the genome-wide average regardless of the focal triplet.

**Figure 3.**
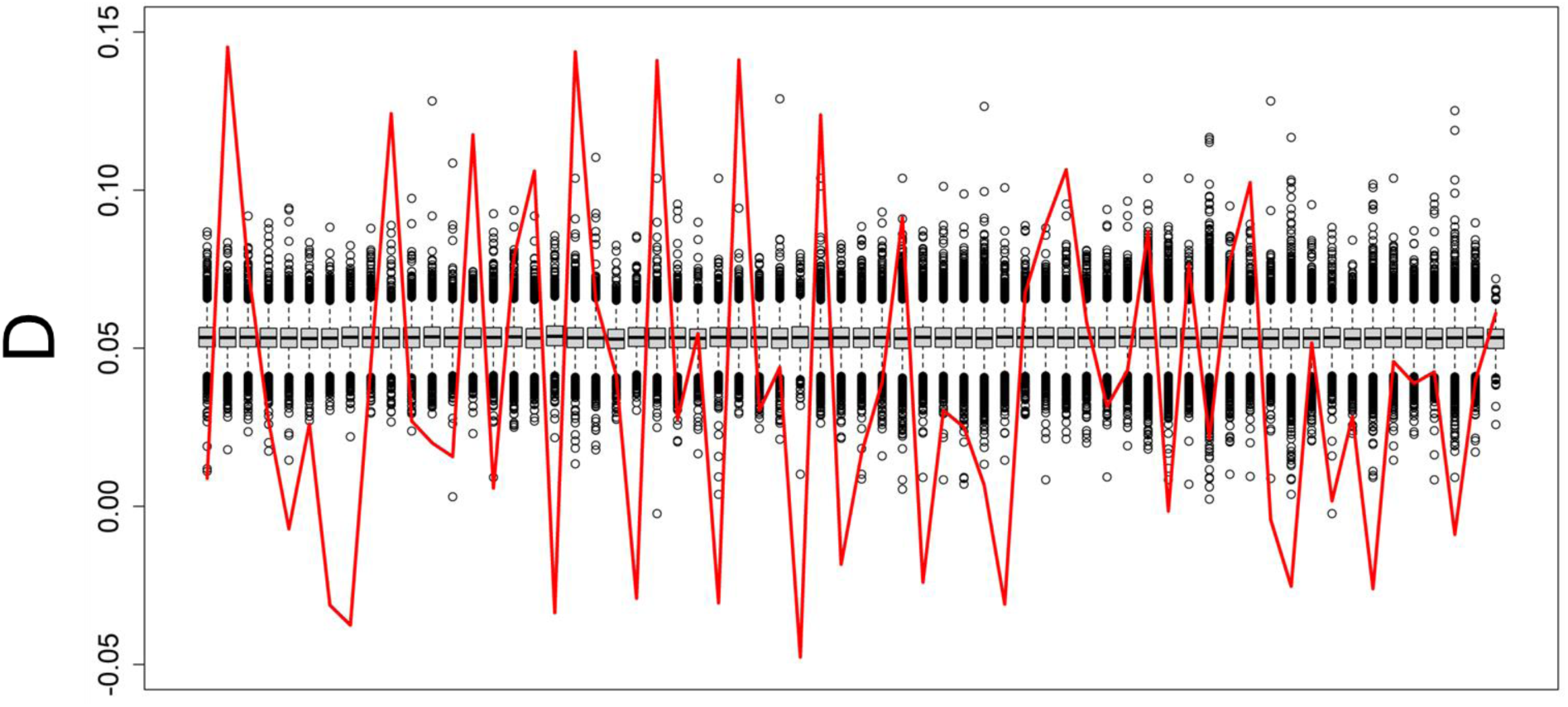
Can triplet-specific variation in D be generated by introgressed fragments? The horizontal axis is triplet identity, with 32 transversions followed by 32 transitions in the usual order. The red line represents genome-wide triplet-specific values for D(Yoruba, French, Altai, chimpanzee). A boxplot is drawn for each triplet, summarizing the distribution of values of expected D (Fig. 2) for all instances on chromosome 1 where that triplet lies at the centre of a valid 40Kb window.

If the variation in D between triplets is driven primarily by introgression, sites most likely generated by introgression should exhibit an exaggerated pattern that is less ‘diluted’ by other sites. We tested this prediction by defining the most likely introgressed sites as biallelic sites where the Neanderthal and chimpanzee alleles differ and where the Neanderthal allele is absent from Africa and occurs at < 50% frequency outside Africa. The overall distribution of such sites along each chromosome is ostensively suggestive of introgression, with most 50kb windows carrying no sites and a small minority of windows (∼0.5%) carrying up to 10 or even 20 sites. However, the frequencies of triplets among introgression-consistent sites are weakly *negatively* correlated with triplet-specific D. The main pattern seen is a massive over-representation of the most mutable XCG transitions (Segurel, Wyman, and Przeworski 2014; Besenbacher et al. 2016): among introgression-consistent sites, XCG transitions make up 64% of all D-informative triplets, while among all other sites they contribute 14.5%. This pattern strongly supports the idea that whatever drives D has little or nothing to do with introgression. Instead, sites carrying alleles not found in Africa appear associated with *de novo* mutations, which, not surprisingly, are disproportionately found at highly mutable sites.

#### Mutation rates are generally higher in Africans compared with non-Africans and mutation rate differences between populations strongly predicts D

To interpret D correctly, it is vital to discover the extent of any variation in mutation rate between human populations. Mallick et al. (2016) estimated that non-Africans have mutation rates that are slightly but significantly higher than Africans. However, their method ignores recurrent mutations by assuming that the chimpanzee is a perfect proxy for the ancestral human allele, and their data have very small sample sizes from each population, which can introduce biases (ESM4). By requiring all three non-human hominins to carry the same allele and using this allele as the presumed human ancestral state regardless of the chimpanzee allele, the number of confounding recurrent mutations should be reduced some 10-fold (chimpanzee->human ancestor ∼ 12 million years, other Hominins -> human ancestor ∼ 1 million years). Applied to the 1KG data, this new measure reveals a modest drop in mutation rate between Africans and non-Africans, with a non-African-to-African ratio of ∼92%. In simulated data, this measure performed well, capturing simulated mutation rate differences where present and revealing rate equality where not. Applied to all pairwise population comparisons in the 1KG data, we see a pattern in which Africans have the highest rates and East Asia the lowest. Given the behaviours of East Asian and South Asian populations, European rates are unexpectedly close to Africa, which may reflect a bias related to the human reference sequence (Fig. 4).

**Figure 4.**
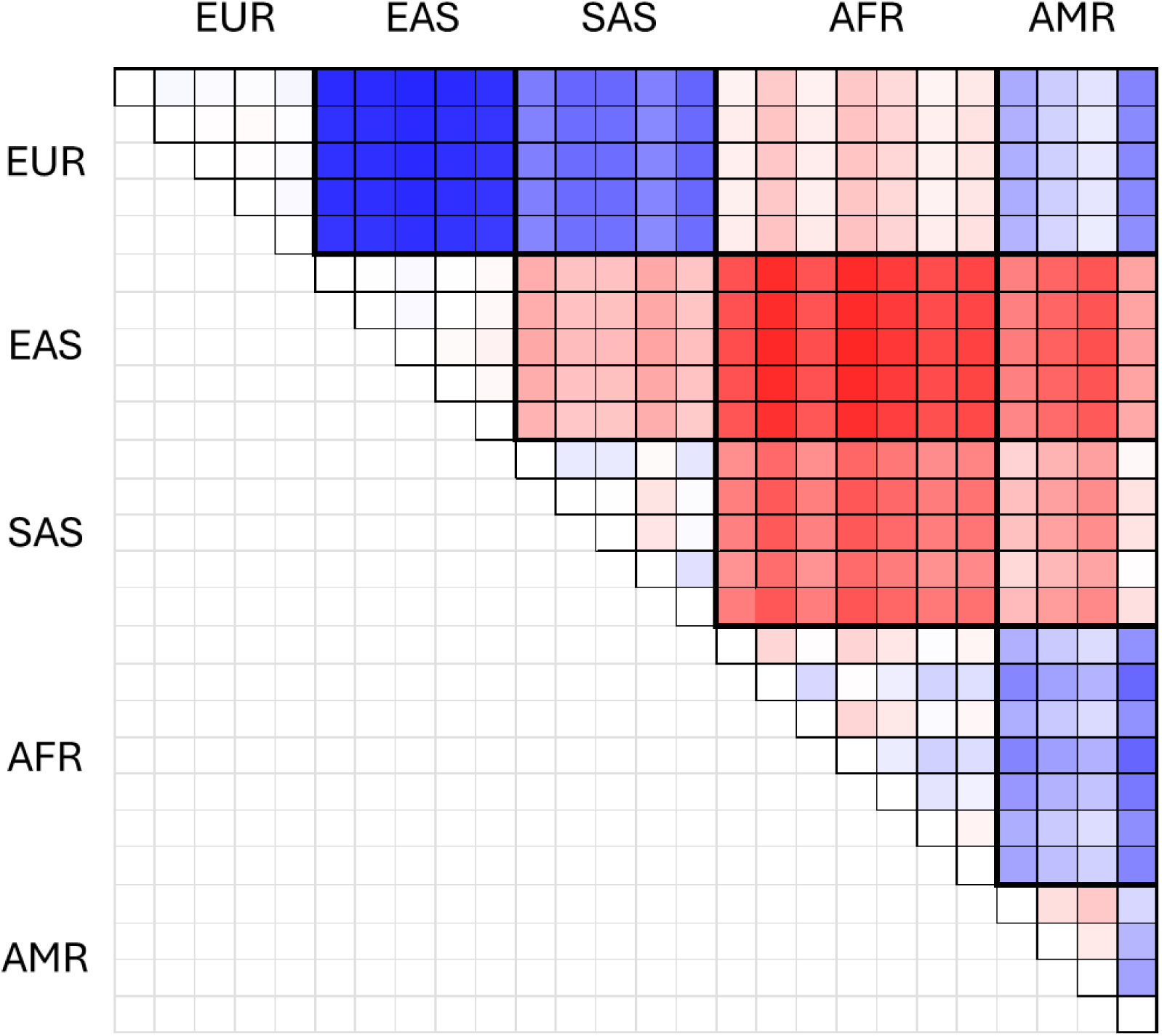
Variation in relative mutation rate between population pairs. Relative mutation rates were calculated using the method of Mallick et al. but replacing the chimpanzee as the assumed ancestral allele in humans with the non-human Hominin allele. Sites where non-human Hominins are polymorphic are excluded because these require one or more mutations outside the human lineage. Ratios vary from 0.94 (darkest red) through parity (white) to 1.06 (darkest blue) and are calculated as row population/column population above the top-left to bottom-right diagonal.

We note that the approach we take here is highly conservative for two reasons: first, our new statistic includes many variants that originated before the out-of-Africa bottleneck, so it underestimates the actual drop in rate that occurred; second, further exploration of the statistic suggests the true relative rate is more likely around 80% (ESM4)

### Testing the distribution and nature of variants that drive positive D

In this section, we examine the relationship between D and allele frequencies in different populations under the two competing hypotheses, an aspect that has received little previous attention.

#### D is dominated by sites where the Neanderthal B allele is fixed outside Africa

If D is driven by introgression, it should be dominated by sites where the Neanderthal B allele is rare outside Africa and absent, or extremely rare inside. We partitioned D according to whether the major allele matches the Neanderthal or the chimpanzee. Absolute values vary somewhat, depending on whether the non-human Hominins are polymorphic or monomorphic and whether the major allele is ‘unambiguous’ (the same inside and outside Africa) or all sites are used (ambiguous sites resolved through an overall majority rule) (ESM 5). Nonetheless, a general pattern is clear where D is strongly positive (0.2 to 0.25) when the human major allele matches the Neanderthal and strongly negative (around -0.15) when the human major allele matches the chimpanzee (Fig. 5). Thus, the classically reported D(African, European, Neanderthal, chimpanzee) of 0.05 represents the difference between two much larger but opposing sub-signals. Crucially, the component that is expected to be driven by introgression, where the Neanderthal allele is the human minor allele, is *negative*, which would classically be interpreted as indicating introgression into Africa.

**Figure 5.**
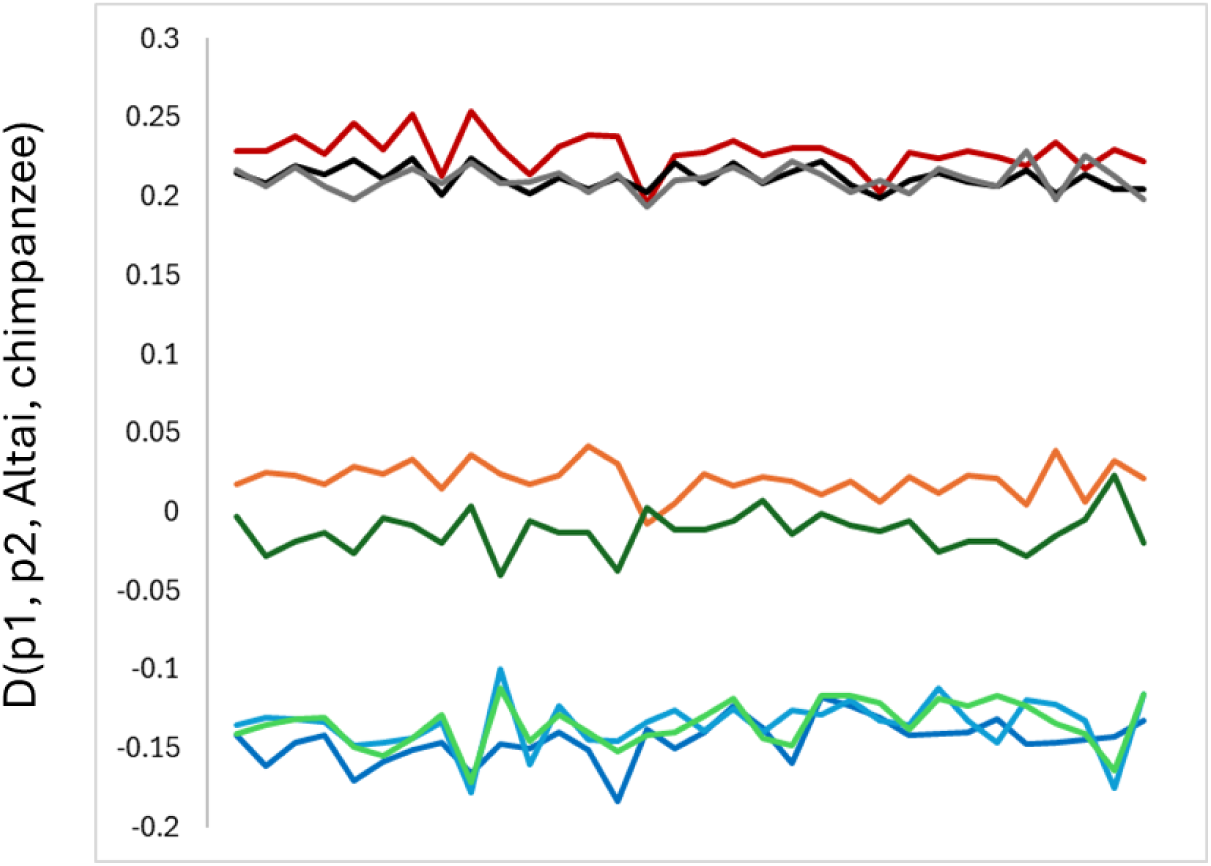
D partitioned according to whether the human major allele matches the Neanderthal or the chimpanzee. Data are for transversions only, and each represents the average of all possible pairwise population comparisons within each regional comparison. For the major allele = Neanderthal component, lines are EAS-AFR (red), EUR-AFR (grey), SAS-AFR (black), EAS-SAS (orange), and for the major allele = chimpanzee component, lines are EAS-AFR (dark blue), EUR-AFR (green), SAS-AFR (light blue), EAS-SAS (dark green).The x-axis is triplet identity(transversions and transitions in the usual order).

In simulated data, similar patterns appear, though they show a strong dependence on the intensity and timing of the bottleneck. In the Gravel model (Gravel et al. 2011; Haller, Ralph, and Messer 2026), D splits into the same positive and negative components seen in real data. However, without a bottleneck, the split disappears, indicating that the out-of-Africa bottleneck dominates the split. Moreover, replacing the Gravel model with a short, sharp step bottleneck prior to introgression actually reverses the pattern, such that sites where the Neanderthal allele is the human minor allele exhibit positive D. Adding introgression to the Gravel model has little impact on the positive component but, as expected, makes the negative component more positive, with higher levels of introgression driving a bigger shift towards being positive. Introducing a mutation rate difference between Africans and non-Africans has very little impact on the negative component but progressively increases the positive component in line with the size of the mutation rate drop (Table S5.1). Thus, the empirical pattern strongly supports a long, slow out-of-Africa bottleneck but arguably tells us less about introgression than we originally thought: the data are entirely compatible with mutation rate variation and argue against introgression at around 1-2%. However, with many uncertainties about the exact form of the bottleneck, this analysis alone does not preclude introgression in the range 0-0.5%. We also plotted the two components on the same axes as the undecomposed D (the original triplet-specific D) and were surprised to find that triplet-specific D exhibits far greater amplitude variation than either sub-component (Fig. S5.1), suggesting some degree of non-additivity.

We next examined how these patterns affect D relative to the expected allele frequencies under the two competing hypotheses. For this, we decomposed D by omitting the normalization step and focusing just on the counts of ABBA minus BABA, which, unlike D, are additive. All sites were classified according to which allele, chimpanzee A or Neanderthal B, is fixed in which population, Africa or Europe, and counted separately for each triplet. We compare the empirical results with simulations under the two hypotheses and summarize them in Fig. 6.

**Figure 6.**
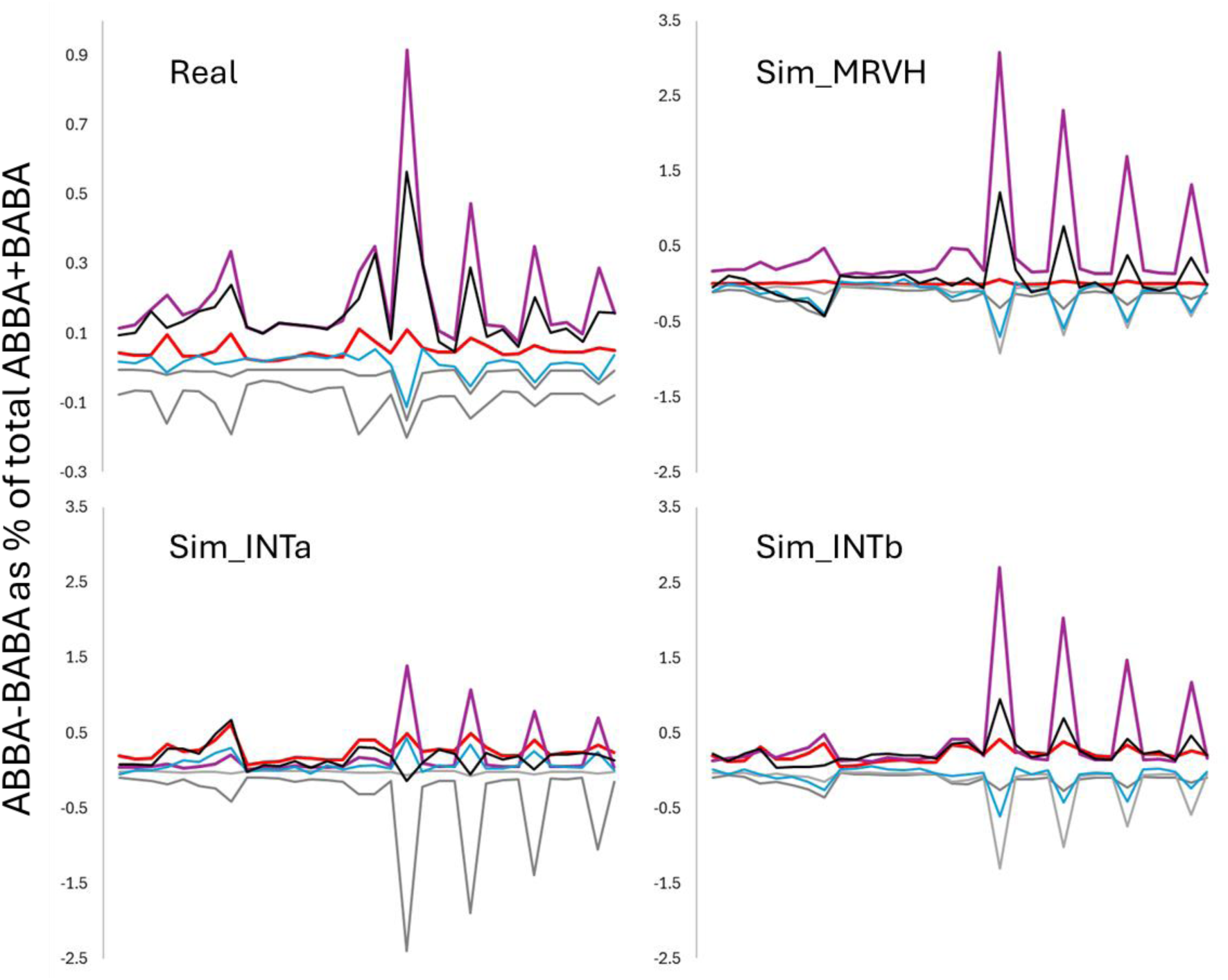
Decomposing D according to which alleles are fixed in which population. ABBAs and BABAs were counted for each triplet (x-axis) in each of five scenarios: A fixed in Africa (red); A fixed in Europe (grey); B fixed in Africa (dark grey); B fixed in Europe (purple); both regions polymorphic (blue) and total counts across scenarios (black). For maximum comparability, all values are expressed as the percent total counts across all triplets and all scenarios. Only transitions are shown because equivalent values for transversions are much lower and less variable. The four panels represent empirical data (Real; note the narrower Y-axis scale), a simulated 75% drop in the non-African mutation rate (Sim_MRVH), 1% introgression with a step bottleneck (Sim_INTa), and 1.5% introgression in the Gravel model (Sim_INTb).

Under the introgression hypothesis, we expect the signal to be dominated by sites where A is fixed in Africa and B is rare in Europe (red), while under the MRVH we expect the signal to be dominated by sites where B is fixed in Europe and A is rare in Africa (purple). In the empirical data (top left panel), the overall signal (black) closely matches the profile from sites expected under the MRVH, with sites expected from introgression contributing very little. We can express the relative proportion of the signal explained by the two hypotheses as *H_1_/H_2_*, the ratio:

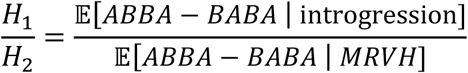

In real data, the ratio = 0.28. In simulations with no introgression but a large drop in non-African mutation rate, the ratio is even more extreme (0.03), confirming next to no contribution from sites expected under introgression. However, in simulations with constant mutation rate and introgression, the ratios are much larger (Sim_INTa ratio = 0.67, Sim_INTb ratio = 1.89). Two additional noticeable trends are apparent. First, in all four scenarios, the profiles are dominated by positive and negative peaks at the four XCG triplets, these peaks being considerably exaggerated in simulated data compared with real data. We do not fully understand why the peaks are so exaggerated, though it is not particularly surprising given that these peaks are driven by recurrent mutations, which scale in frequency with mutation rate squared. Second, Sim_INTa and Sim_INTb show large differences in the relative sizes of the XCG-positive and XCG-negative peaks, despite having nominally equivalent demographies, with both experiencing a bottleneck that caused around a 25% reduction in heterozygosity. The reason relates to the timing and length of the bottleneck. In INTa, a short, sharp bottleneck occurs immediately before introgression, whereas in INTb, which uses the Gravel model, the bottleneck is extended, so introgression occurs near the start, and introgressed haplotypes have more opportunity to be influenced by increased drift.

We observe much higher XCG peaks in simulated data than in empirical data, suggesting that recurrent mutations have an exaggerated impact. One possible explanation relates to the lability of mutation hotspots. It has been estimated that chimpanzees share few, if any, recombination hotspots with humans (Ptak et al. 2005), and that recombination and mutation rates tend to be correlated (Lercher and Hurst 2002; Halldorsson et al. 2019). Although not modeled, it seems reasonable to suggest that widespread changes in hotspot locations could considerably reduce the number of recurrent mutations relative to the mean mutation rate.

### Testing a possible mechanism capable of generating the observed patterns

The introgression hypothesis’s general failure to fit the data implies a large mutation-rate difference between Africans and non-Africans, raising the important question of the underlying mechanism (ESM6). Variation in mutation rate might be caused by (as yet unknown) variants in enzymes involved in DNA replication (Liu and Samee 2023). Another possibility is that heterozygosity modulates mutation rate, such that the large loss of diversity during and after the out-of-Africa event (Prugnolle, Manica, and Balloux 2005) caused a parallel reduction in mutation rate (Amos 2013; Yang et al. 2015; Amos 2016). Because this mechanism is arguably the only currently proposed explanation for the mutation-rate gradient we document across Eurasia, we next tested the hypothesis that variation in heterozygosity drives variation in mutation rates, which, in turn, drives D.

#### Heterozygosity difference predicts relative mutation rate across pairwise population combinations and across the genome

To assess whether heterozygosity differences between populations predict corresponding differences in mutation rate, we estimated relative mutation rates between all pairwise population comparisons, using an adapted form of the method of Mallick et al. (2016) (ESM4), where the chimpanzee is replaced as the outgroup by the consensus of all three archaics. For this analysis, we require all non-humans to carry the same allele, thereby completely decoupling relative mutation rate from D (each site can contribute to only one measure, never both). When plotted against heterozygosity difference, we find ubiquitous and highly significant relationships both overall and within each major geographic region (Fig. 7, Fig. S4.5). Equivalent plots based on all sites, regardless of the chimpanzee state, are extremely similar. Interestingly, many of the slopes also differ significantly from zero, as expected under a model in which changes in heterozygosity drive changes in mutation rate. This is because admixed populations like those in America have experienced potentially large changes in heterozygosity over a short timescale that would allow little time for any downstream change in mutation rate to become manifest. We have already shown that mutation-rate differences and D are correlated, so it is no surprise that heterozygosity differences also predict D.

**Figure 7.**
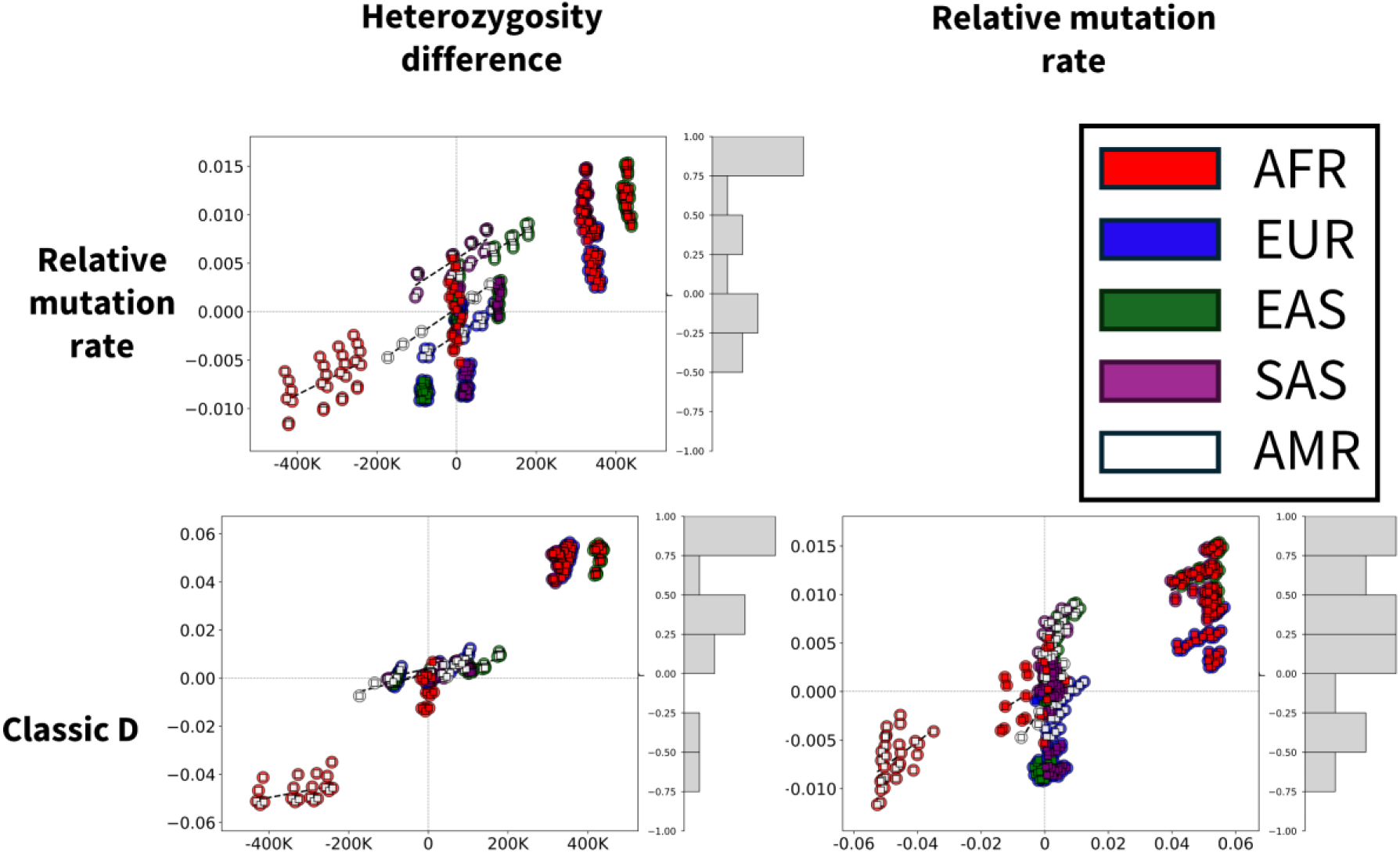
Relationships among heterozygosity difference, the D statistic, and relative mutation rate. Pairwise scatterplots compare heterozygosity difference, the classic D statistic, and the relative mutation-rate statistic across 1KG populations. Each point represents a population pair and is coloured by geographic region. Most regressions (dashed lines) are positive (see histograms for r distribution). Although relative mutation rate is usually expressed as MU_Eur / MU_Afr, for these plots we used a statistic that is directly analogous to D: (MU_Pop1 – MU_Pop2) / (MU_Pop1 – MU_Pop2), where Pop1 = Africa / the population closest to Africa. Positive values therefore indicate a higher rate in Africa. Note, all regressions involving relative mutation rate become far stronger if we exclude sites where all non-humans carry the alternate human allele (ESM 8).

Across the genome, natural selection has modulated the rate at which heterozygosity was lost out of Africa, with positive selection outside Africa sometimes acting to accelerate loss of diversity and balancing selection sometimes acting to retard loss or even increase heterozygosity (Amos and Bryant 2011). We therefore repeated the above analysis for non-overlapping 50kb windows across the genome, using pooled African and pooled non-African samples, carefully matched for sample size. Trends reported for whole-genome comparisons are also seen across the genome, where variation in heterozygosity difference is also a potent predictor of relative mutation rate and D. We tested a prediction of MRVH, that the relationship between D and mutation rate will depend on the states carried by the outgroups, as illustrated in Fig. 1A. At PBBBA sites, a higher rate of recurrent B->A mutations in Africa will drive an excess of ABBAs. In contrast, at PBBAA sites, where B is derived in Neanderthals and the human ancestral state is A, recurrent mutations in humans will be A->B, and if these occur at a higher rate in Africa, they will drive an excess of ABBAs. We find clear evidence for this: D is positively correlated with relative mutation rate at PBBBA sites but negatively correlated at PBBAA sites (ESM3, Fig. S3.3, Fig. S3.4 left-hand panel). This is the antithesis of expectations under the introgression hypothesis, where D should be independent of mutation-rate variation and should not vary depending on whether the Denisovan allele is the same or different. The change in slope direction appears to pose an insurmountable challenge to explanations based on either artifacts or introgression. Moreover, we find that differences in heterozygosity strongly predict relative mutation rate, with higher heterozygosity in Africa predicting a higher mutation rate in Africans (Fig. S3.4 right-hand panel).

For comparison, we also simulated introgression (Fig. 8). To ensure any trends would be apparent, even if weak, we deliberately used a higher rate than might have occurred in humans: 1.5%, giving an overall D of 0.11, about double the empirical value. We find a very clear pattern: both graphs have effectively two sections. Most windows are not affected by introgressed material and show positive heterozygosity differences reflecting the out-of-Africa bottleneck. In these sections, both D and relative mutation rate show no relationship with heterozygosity difference. In contrast, in windows with introgressed material, non-African heterozygosity increases, yielding negative heterozygosity differences. In these windows, the introgressed material increases D and decreases relative mutation rate. The mutation-rate reduction occurs because we used the Neanderthal allele as the assumed human ancestral allele, as we do when analysing empirical data, and windows with Neanderthal material inevitably appear closer to Neanderthals. Thus, mutation-rate differences do not create artefactual D or signals of increased or decreased mutation rate; this happens only when introgressed material is present, and here the trends run opposite to those seen in empirical data.

**Figure 8.**
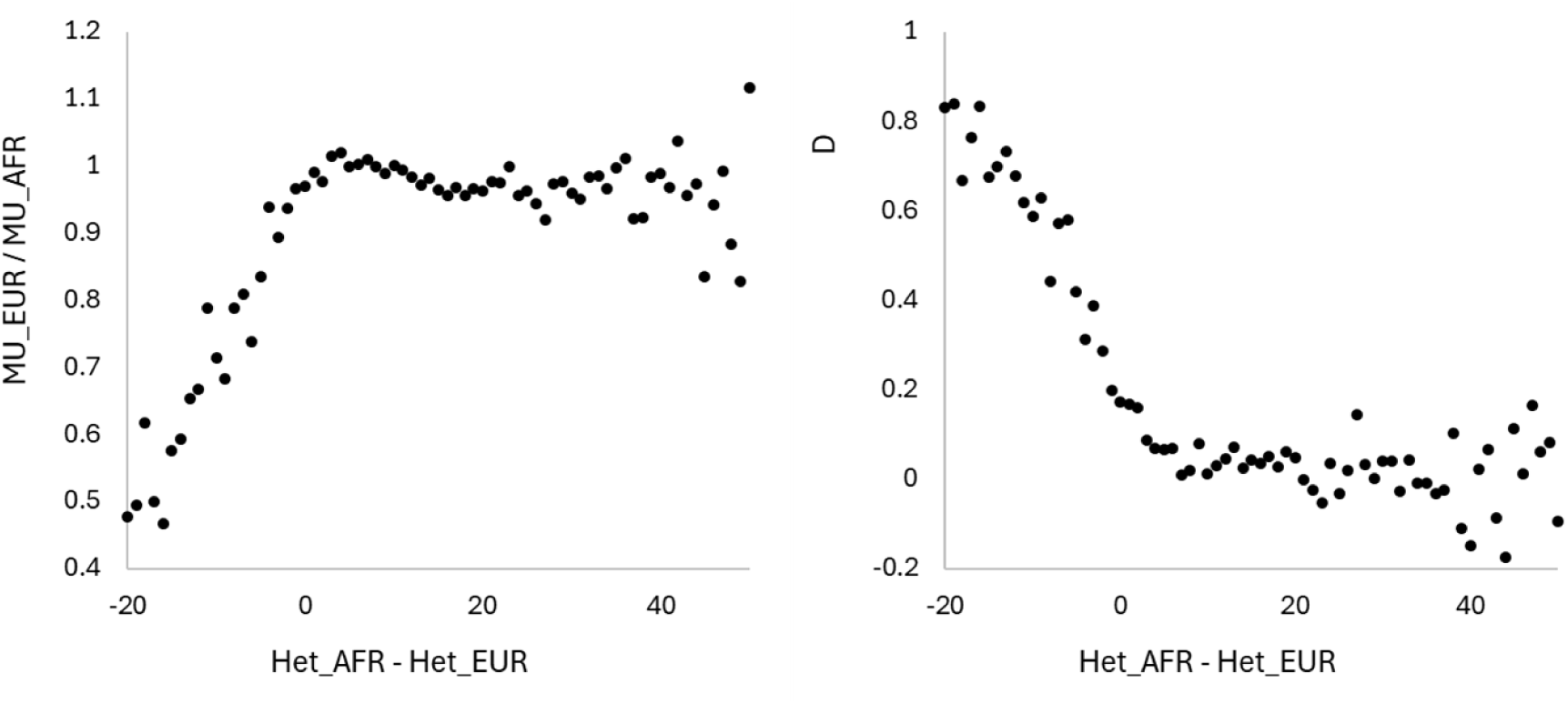
Relationships between heterozygosity difference, relative mutation rate and D under simulated introgression. Heterozygosity was summed in each population within each 50kb genomic window, and the bins (x-axis) are the integerised values. Introgression was simulated at a rate of 1.5%, giving an overall D of 0.11. Each plot represents 500 simulation runs, each of a different 0.5Mb fragment.

Based on these findings, we can reconstruct a plausible sequence of events underlying the observed D signal (Fig. 9). Before the out-of-Africa event, Africans and the ancestors of non-Africans experienced broadly similar mutation processes, with ILS-generated ABBA and BABA configurations occurring at comparable frequencies. The subsequent loss of heterozygosity in non-Africans was accompanied by a reduced mutation rate, creating population differences in the frequency and spectrum of recurrent mutations. As heterozygosity and mutation rates gradually recovered in non-Africans, these differences diminished. In this framework, population-specific recurrent mutations acting on different ancestral configurations generate opposing components of D, and their imbalance produces the observed positive net D. By contrast, components expected from Neanderthal introgression (had they existed) contributed little to the overall signal. The collective evidence provides a coherent reconstruction linking demographic history, heterozygosity, mutation-rate variation, and recurrent mutation to the emergence of D.

**Figure 9.**
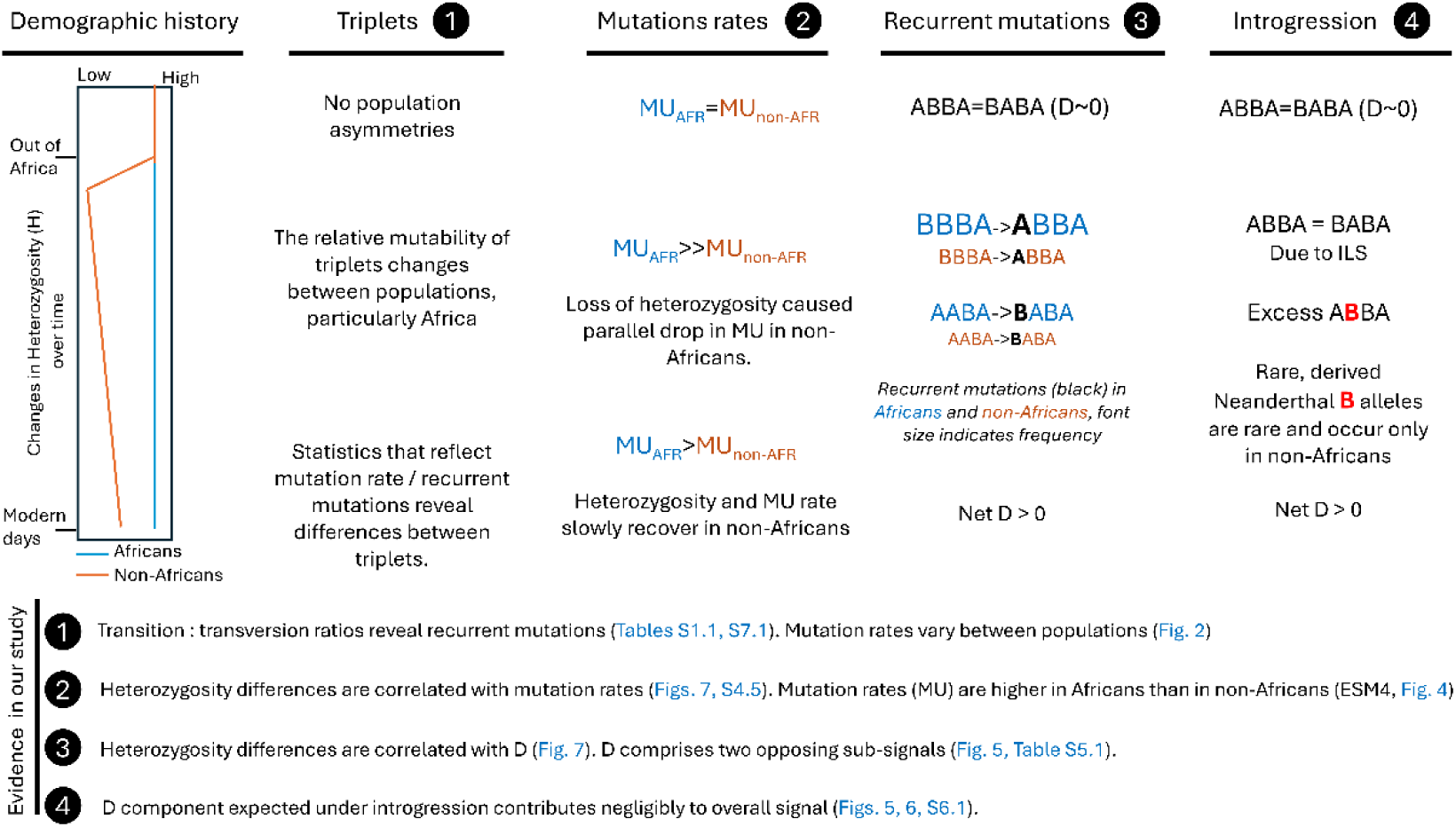
A reconstruction of the changes during human evolution in light of the evidence. The schematic follows African and non-African lineages from before the out-of-Africa event to the present, tracking changes in triplets, mutation rates, D, and introgression. ABBA and BABA configurations illustrate how recurrent mutations can modify sites originally generated by ILS. Events are cross-referenced with the supporting evidence.

## Discussion

The D statistic, also known as the ABBA-BABA test, was proposed over a decade and a half ago (Green et al.) as a tool to infer archaic introgression into humans and has since been used extensively (∼3000 citations) to detect interbreeding or admixture. A positive D indicating that Neanderthals share more bases with non-Africans than with Africans was widely interpreted as evidence for introgression. Here we scrutinise D by examining the assumptions on which it is founded. We show that both assumptions are false. Then, for the first time, we consider side by side both possible explanations for elevated similarity between non-Africans and Neanderthals: introgression reducing the Non-African-Neanderthal distance versus a relatively higher mutation rate in Africans increasing the African-Neanderthal distance (Fig. 1B). At every step, we compare the patterns seen in real data with the output of realistic simulations. We find unambiguous support that mutation-rate differences are the primary, and probably only, driver of positive D. Thus, while interbreeding likely did happen, any resulting introgression was negligible.

The most important flaw we expose is that the assumption that recurrent mutations are so rare they can be safely ignored does not hold (Green et al. 2010; Durand et al. 2011; Sankararaman et al. 2014; Sankararaman et al. 2016). We suspect this assumption originates from naïve calculations based on a genome-wide average mutation rate of ∼10^−8^ per site per generation, a rate that generates virtually no recurrent mutations. However, this average rate hides massive underlying variation (Aggarwala and Voight 2016; Koppetsch, Malinsky, and Matschiner 2024), with up to 100-fold variation in rate along a chromosome and rates varying across the mutation spectrum by a factor of 40. We do not know whether these two sources of variation are additive or interact in a more complicated way. However, because the probability of a recurrent mutation scales with mutation rate squared, the most mutable sites plausibly generate recurrent mutations up to 10,000 times more often than expected from the average rate. We show this in both real datasets used to study possible introgression and in realistic simulated data: about 1-3% of polymorphic sites have experienced more than one mutation. 1-3% may appear modest, but it far exceeds the fraction of sites that typically carry the signal used to infer introgression. Recurrently mutated sites outnumber D-informative sites by roughly two orders of magnitude, rendering the assumption that recurrent mutations occur at a negligible rate untenable.

Throughout our study, we show time and again how recurrent mutations distort and undermine the validity of apparently reasonable approximations, nowhere more so than in estimating mutation rates. When Mallick et al. (2016) counted derived variants on human lineages assuming the chimpanzee provides an excellent proxy for the human ancestral state, they concluded that Africans have a slightly but significantly lower mutation rate than non-Africans. However, recurrent mutations create an inverted signal because mutations between chimpanzees and humans mean the assumed and true human ancestral states differ, causing the wrong lineage to be credited with a derived variant. Although no perfect solution exists, we showed that the non-African mutation rate is likely around 10-20% lower than the African rate. Moreover, this estimate necessarily relies on all variants (the method is robust to differences in demographic history only if all variants are analysed regardless of frequency), and most common variants predate the African–non-African split. Consequently, the figure of 10-20% would be appreciably larger if it were based only on variants arising from mutations that occurred *after* the non-African lineage left Africa (Table S4.1).

We next used realistic forward simulations to ask whether a plausible drop in the non-African rate could be sufficient to drive realistic values of D. We find that the 10-20% drop in our measure of relative rate, calculated over all sites, could equate to a 50-70% drop in actual rate since the African–non-African split (Tables S4.1, S7.1). Furthermore, this rate drop generates D values consistent with empirical values from real data. Unfortunately, it is difficult to be more precise. We do not know the true human ancestral state, nor do we have good estimates for how every element of the mutation spectrum varies between human populations (Harris 2015; Harris and Pritchard 2017). Uncertainties also remain about how rate variation across the mutation spectrum interacts with variation in overall mutation rate along a chromosome, through hotspots and coldspots (Aggarwala and Voight 2016; Koppetsch, Malinsky, and Matschiner 2024). Moreover, if mutation rate in humans has generally increased over the last few thousand generations, pushing a greater proportion of variant origins towards the present, a much smaller drop in the non-African rate would be required to produce a given value of D. At present, we struggle to distinguish between a modest drop in mutation rate out of Africa coupled with a generally increasing rate in humans or a larger drop in rate and no general increase (ESM 4). However, our analyses appear to eliminate introgression as anything but a minor driver of D (at most), leaving variation in mutation rate as the only viable explanation. Finally, we know that many mutations have occurred between humans and chimpanzees. Consequently, the approximate parity of rates reported by Mallick et al. must include both a component where the relative rates are estimated correctly and a component where the signal has been inverted; we must subtract the latter from one side and add it to the other. Ironically, estimated parity shows that an imbalance must exist and, while the exact size is difficult to estimate, recurrent mutations are disproportionately more likely at the fastest-evolving sites, so the actual imbalance is likely larger than intuition might suggest from the average level of human–chimpanzee divergence. An important consequence of these simulations is that greater variation in mutation rate across sites generates more recurrent mutations for a given average mutation rate. Thus, modelling the full triplet mutation spectrum substantially reduces the mutation-rate difference required to generate realistic values of D. Because our simulations do not include additional regional variation such as mutation hotspots, they are likely conservative.

The key observation that D varies strongly when calculated separately for each triplet motif across the mutation spectrum further supports strong mutation-rate variation. Introgression theory indicates that such pronounced triplet-specific variation should not occur if mutation rates do not vary between populations, but it is consistent with the works of Harris (Harris 2015; Harris and Pritchard 2017), who showed that mutation rates of individual triplets vary globally, particularly between Africans and non-Africans. However, the implications go much further. All informative sites on an introgressed fragment must contribute equally to D for that fragment. As a result, as long as introgressed haplotypes carry several, more or less random triplets, it is effectively impossible for different triplets to give different values of D. We confirm this expectation by showing how, in real data, the observed variation in D across triplets is completely removed when the data are analysed as introgressed fragments. Specifically, we show that a typical 40kb fragment contains enough D-informative triplets that the expected D of each fragment lies close to the genome-wide value of 0.05. Furthermore, inter-triplet variability in D is far greater in CpG islands. We are forced to conclude that the variation in triplet-specific D seen in real data precludes introgression from contributing more than a minor amount to D.

Further evidence against introgressed material contributing appreciably to D comes from comparisons of triplet-specific D between non-African populations/regions. These also reveal large variation that parallels the trends seen in African–non-African comparisons, with the same triplets tending to give high and low values. Under the introgression hypothesis, different triplets can give different values of D only if introgressed haplotypes are favored or purged depending on the triplets they carry. This is biologically implausible for many reasons, not least the fact that selection acts on base substitutions and not on the identity of the two flanking bases. Nonetheless, even if selection could act on haplotypes based on the triplets they carry, the fact that all non-African comparisons have D profiles that fluctuate both above and below zero stretches credibility even further. Such a pattern would require selection to act both positively and negatively on the same triplet in different parts of Eurasia.

We also note more subtle trends that argue against introgression. In Fig. 2, the regional comparison with the largest overall D, Africa–East Asia (purple), dominates not only the highest peaks, but also tends to be visible at the lowest troughs. In other words, higher D is associated with more extreme variation across the mutation spectrum, not merely a vertical upward shift for all triplets as would be expected if the pattern were driven by introgression. More generally, even if introgression did create variation in D across triplets, variation in the amount of introgressed material would manifest as simple y-axis shifts of the same basic profile. In reality, it is easy to find instances where a trough in one profile corresponds to a peak in another. Such ‘conflicts’ seriously challenge explanations built around introgression.

Surprisingly, triplet-specific D also varies in simulated data even when mutation rates do not vary geographically (though they do differ between triplets). This appears to contradict theory, where the mutation rate term drops out as long as mutation rate does not vary between populations, conditions that in the simulated data hold for each triplet treated separately. Although we lack a full explanation, the correlation between triplet-specific D in simulations and empirical values argues strongly that this is a real emergent property. We speculate that this may reflect the way more mutable triplets become locally rarer in mutation hotspots as they mutate to other states, but more work is needed. Crucially, when we introduce introgression to the simulations, triplet-specific D no longer correlates even weakly with empirical values. This shows that any inter-triplet variation in D produced by introgression differs radically from the empirical pattern, implying that processes other than introgression drive the observed inter-triplet variation in human genomes.

The finding that D varies massively across the mutation spectrum suggests that D is driven mainly or entirely by a mechanism operating through individual bases, not introgressed fragments. Mutation rate variation is the obvious candidate. Mutation rate, as estimated by our adjusted version of Mallick et al.’s method, correlates strongly with D across the globe, both between African–non-African population pairs and between African and non-African pairs. Thus, despite being calculated on two entirely non-overlapping sets of sites, the two measures appear to be closely and ubiquitously linked. Such a pattern is neither expected from a model based on introgression, not least because the current model of introgression assumes that mutation rate does not vary, nor is it seen in data simulating introgression. Instead, the correlated measures confirm that both reflect the same underlying property: relative mutation rate. This should come as no great surprise because the two measures are identical in the way they are calculated with respect to humans and differ only in the conditioning applied to the outgroups: for relative mutation rate we require the chimpanzee and all non-human Hominins to carry the same allele, while for D the chimpanzee and Altai must differ. Nonetheless, the correlation helps tie D in humans to variation in mutation rate rather than to introgressed material.

An important question is why mutation rate should vary as our analyses suggest. We address this by extending a previous idea (Amos 2013) that heterozygosity modulates mutation rate, with more and different types of mutation occurring in regions with higher heterozygosity. Fit to this model appears excellent, well beyond showing that the big drop in heterozygosity that occurred post ‘out of Africa’ (Prugnolle, Manica, and Balloux 2005) led to a corresponding drop in relative mutation rate. First, the progressive drop in heterozygosity (Prugnolle, Manica, and Balloux 2005) moving eastwards across Eurasia fits with a parallel drop in mutation rate and an associated increase in D (Wall et al. 2013). Second, more generally, wherever differences in heterozygosity are seen, so too are differences in mutation rate and correlated variation in D, regardless of where the population sample comes from, including comparisons within Africa. Third, the out-of-Africa loss of diversity varies greatly across the genome, and this again generates a strong correlation between heterozygosity loss and relative mutation rate.

These patterns explain not only why introgression seems to be inferred essentially wherever it is looked for, but also why so many links are found between genes under strong selection (Sankararaman et al. 2014; Gittelman et al. 2016; Reilly et al. 2022; Iasi et al. 2024), specifically in Asia (Kanis et al. 2026), and supposed archaic origin: Strong selection appears to impact local mutation rates (Monroe et al. 2022) and we suggest a simple mechanism whereby selection impacts heterozygosity which in turn drives changes in mutation rate. Finally, we often hear the argument that D is imperfect, so evidence such as we present here does not affect the introgression narrative because other, stronger evidence comes from, for example, introgressed haplotypes (Huerta-Sánchez et al. 2014; Skov et al. 2020; Reilly et al. 2022). However, this is misleading. D is not flawed for failing to capture an introgression signal: if a signal is present, D will certainly capture it. The key flaw lies in interpreting D. Specifically, it rests on the near-ubiquitous assumption that the signal D captures is unambiguously due to introgression rather than other evolutionary processes. Evidence from introgressed haplotypes may appear strong, but the methods used to infer them rely heavily on the same flawed assumptions that underpin D, so they need revisiting. Indeed, we show here that a model based on fragments cannot be reconciled with the level of variation in D seen between triplets. More generally, a model not challenged against a meaningful null model is assumed rather than demonstrated. To be considered rigorous, any test of introgression must strive to compare all possible explanatory hypotheses side-by-side and without favour, regardless of how likely they are considered *a priori*. Previous studies acknowledge that variable mutation rates *could* drive D, but exclude this model by assuming a constant mutation rate and that recurrent mutations are negligibly rare. As we show, these assumptions are both false. We urge future studies to always seek tests that properly compare predictions from mutation rate variation and introgression. When this is done, we are confident that support for the introgression narrative will evaporate.

As with any study, our analyses have some limitations. We acknowledge that residual errors are unavoidable in ancient DNA. However, by using all available high-quality archaic genomes and conservative consensus-based filtering, we expect any remaining biases to be minimal and equally applicable to both hypotheses. Moreover, forward simulations and empirical observations independently support our principal conclusions.

Our study helps resolve several long-standing conundrums and issues with the introgression narrative. First, no one has yet discovered Neanderthal mitochondrial DNA or Y chromosomes in humans (Petr et al. 2020). Although some modelling suggests this is not incompatible with interbreeding leading to introgression (Currat and Excoffier 2011; Petr et al. 2020), the strong negative selection required would imply a considerable fitness barrier to the spread of introgressed material. Second, D increases from west to east across Eurasia (Wall et al. 2013; Amos 2021), a pattern that is extremely difficult to explain with a simple introgression model in which Neanderthals lived mainly in the west. Third, the introgression hypothesis predicts that almost the entire signal should be carried by heterozygous sites in non-Africans, but the exact opposite is found: the signal is carried almost entirely by heterozygous sites in Africans (Amos 2020). Fourth, a major inconsistency appears between the inferred diversity of archaic hominin lineages, which implies repeated speciation events, and evolutionary theory, which holds that even modest gene flow strongly inhibits speciation (Barton and Bengtsson 1986; Slatkin 1987). Yet, under the current paradigm, researchers infer interbreeding and introgression for nearly every encounter between hominin groups and frequently invoke additional, undescribed “ghost” lineages to reconcile discrepancies in the genetic data (e.g., Durvasula and Sankararaman 2020; Gopalan et al. 2021; Pfennig et al. 2023; Zhang et al. 2026). Although such unsampled populations may of course have existed, invoking an unobserved lineage to explain observations that do not fit a preferred model risks becoming unfalsifiable: methodologically, a “ghost” introduced into a genetic model solely to account for otherwise unexplained patterns is little different from invoking a ghost to explain an otherwise unexplained natural phenomenon: an unobserved entity should not acquire explanatory force merely because it makes the observations fit the preferred model. Ignoring population structure in hominin evolutionary modells can also lead to interbreeding inferences (Tournebize and Chikhi 2024). All these issues are resolved if introgression, as opposed to occasional interbreeding, was minimal or absent and instead if D is driven by the evolution of mutation rates across the mutation spectrum, modulated by the large changes in heterozygosity that are either documented during the out-of-Africa event and subsequent dispersal across the world, or seem likely in the small populations that likely characterized early Hominin evolution. Ultimately, the goal is not to defend a particular narrative of human evolution, but to identify the model that best explains the evidence. If a mutation-rate framework accounts for the observations more parsimoniously and with fewer auxiliary assumptions than pervasive introgression, then it deserves to be taken seriously as the leading explanation.

## Conclusions

Our analyses consistently favour the Mutation Rate Variation Hypothesis over archaic introgression. Dominated by recurrent mutations, the D statistic varies strongly with triplet context and is driven by allele-frequency patterns inconsistent with introgression but predicted by mutation-rate variation. Although introgression may appear more parsimonious because a single exchange event can explain a shared allele, this intuition is misleading: recurrent mutation is a ubiquitous feature of genome evolution, whereas introgression is not uniquely required to explain the observed patterns. Taken together, our results suggest that mutation-rate variation provides a more parsimonious and complete explanation of the D statistic, with any contribution from Neanderthal introgression likely negligible. This conclusion does not depend on fitting a single statistic. The same mutation-rate model simultaneously accounts for empirical mutation-rate estimates, the overall value of D, the enrichment of recurrent mutations, and the opposing allele-frequency components that dominate the D signal.

## Methods

### Dataset

The HGDP deep-resolution dataset (Bergström et al. 2020) on GR38 aligned to the high-coverage Vindija Neanderthal (Prüfer et al. 2017), Altai Neanderthal (Prüfer et al. 2014) and Denisovan (Meyer et al. 2012) were kindly provided by Anders Bergström. Although we opted not to use the HGDP data in the analyses presented here, we are keen to facilitate maximally comparable parallel analyses in the future and used the archaic Hominin alignments as the basis of a set of master files we generated, summarising the high coverage 1000 Genomes data (1KG) as a series of chromosome-specific master files each containing the fields: 1 = chromosome, 2 = location on GR38, 3 = reference allele; 4 = alternate allele; 5-30 = alternate allele frequencies in each of the 26 populations (see order below); 31-32 = alternate allele frequencies in African and outside Africa; 33 = major allele 1; 34 = major allele 2; 35 = triplet; 36 = non-human allele codes. Populations are always ordered as they appear in the VCF files: 5 x Europe (GBR, FIN, CEU, IBS, TSI); 5 x East Asia (CHB, CHS, CDX, KHV, JPT); 5 x South Asia (GIH, STU, ITU, PJL, BEB); 7 x Africa (LWK, ESN, MSL, GWD, YRI, ASW, ACB); and 4 x America (MXL, CLM, PUR, PEL). All population frequencies are based on their first 50 individuals (= 100 chromosomes) encountered in the original phased VCF file downloaded from http://ftp.1000genomes.ebi.ac.uk/vol1/ftp/data_collections/1000G_2504_high_coverage/working/20201028_3202_raw_GT_with_annot/. Thus, if individuals were numbered according to their population origin and the order they appear in the 1KG data, each population would be represented by sample numbers 1:50. African and non-African frequencies are based on the first 84 individuala = 168 chromosomes encountered from each of the five non-admixed African populations (i.e., excluding ASW and ACB) and 28 individuals = 56 chromosomes from each of the five populations in Europe, East Asia and South Asia (i.e., all but America), giving the same total number of individuals/chromosomes sampled in each category. We use two definitions of the major allele. Major allele 1 is referred to as ‘unambiguous’ and is based only on sites where the same allele is major both inside and outside Africa (reference allele = 0, alternate allele = 1, sites where African and non-African major alleles differ = -1). Major allele 2 was coded the same way, but there are no -1 calls because, at sites where the major allele differs between regions, samples are pooled, and the major allele is the one with the higher overall frequency (ESM8). Triplet is the inferred ancestral mutating triplet under the assumption that Major allele 2 is the ancestral human allele. Triplets were coded as: 0-31 as transversions outside CpG islands, 32-63 as transitions outside CpG islands, 64-95 as transversions inside CpG islands and 96-127 as transitions inside CpG islands. CpG islands are defined as regions where windows of 200bp have GC content > 50% and an observed minus expected number of CpG motifs of 0.6. Non-human codes are as follows: 0 = X1X0; 1=X0X1; 2=0000; 3=1111; 4=0001; 5=1110; 6=1100; 7=0011, where the taxa are arranged: Vindija, Altai, Denisovan, chimpanzee and 0 = reference allele, 1 = alternate allele, X = any. All categories are non-overlapping such that category 6 is not a sub-category of category 0. Instead, to obtain all sites where Altai = 1 and chimpanzee = 0, one would select categories 0, 5 and 6.

### Data Analysis

Data were processed using a number of custom scripts in C++. The analyses involve calculating D in a variety of different ways, depending on the alleles carried by the outgroups, the human major allele, and which triplet mutates. As such, the input files we provide can be easily processed using scripts in any language. We include an example C++ script that analyses 50kb windows for African–non-African heterozygosity differences and predicts various forms of D for three pairwise regional comparisons, which should be easy to adapt to generate other outputs.

### Estimation of relative mutation rate and D

Relative mutation rate can be assessed by counting derived changes on each lineage in a pair, and when based on all informative sites, it is robust to differences in demographic history. However, the human ancestral state is unknown, and the chimpanzee state used by Mallick et al. is a poor proxy for the human ancestral state because of the frequency of recurrent mutations (see text). We explored several alternatives based on non-human Hominins and the human major allele (ESM4) and found that the best option is to use sites where the Altai, Vindija, and Denisovan all carry the same allele, and assume this is also the human ancestral state. We consider two versions. When the chimpanzee allele is ignored (sites of form PBBBX), the number of sites included is maximised but the moderately small subset where the chimpanzee carries a different base (PBBBA) contributes to D and they are enriched for sites carrying recurrent mutations. The second version excludes these sites (uses only PAAAA) and has the benefit of removing possible noise and completely decoupling estimates of D from estimates of relative mutation rate. On the downside, this second version risks reducing robustness to differences in demographic history.

Having decided on the major allele, relative mutation rates (RMR), were calculated probabilistically in a manner exactly analogous to the methods used to calculate D:

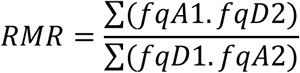

Where fq = frequency, A = the inferred ancestral allele (i.e., the major allele), D = the inferred derived allele, and the numbers indicate the two populations being compared. As written, RMR is the mutation rate in population two relative to the mutation rate in population one.

D is calculated probabilistically in a directly analogous fashion. At any given site where the Altai (B) and chimpanzee (A) bases differ, we calculate two probabilities: (i) of drawing a B allele from population 1 and an A allele from population 2; and (ii) of drawing an A allele from population 1 and a B allele from population 2. These quantities are equivalent to the top line and the bottom line in the equation above after replacing ancestral and derived with matching the Neanderthal and matching the chimpanzee, and represent the probabilities of drawing a single allele from each population and observing a BABA or an ABBA. These probabilities are then summed across sites to give the expected numbers of BABAs and ABBAs for a comparison between two haploid genomes. As usual, D is calculated as the normalised difference between the two: (ABBA – BABA) / (ABBA + BABA).

### Counting recurrent mutations and transition-transversion ratios

To minimize the impact of sequencing or alignment errors, we conservatively required that each allele be present in three or more copies across the 1KG data, without restrictions on geographic distribution. Recurrent mutations were deemed to appear as SNPs with three or four different alleles. Such sites must have undergone at least two mutations in their ancestry, one of which must have been a transversion. Since transition rates are approximately twice those of transversions, recurrent transition events that restore a previously existing allele are effectively “silent” and do not generate additional observable alleles. Consequently, tri- and tetra-allelic sites represent a conservative lower bound on the true frequency of recurrent mutations.

### Forward simulations

To explore the impact of variable mutation rates, we used the forward simulation package SliM v5.1 (Haller, Ralph, and Messer 2026), which allows modelling of elements that are difficult or impossible to model using a coalescent framework. Our basic population model was based on the recipe of Gravel et al. as listed in the SLiM manual (the ‘main’ simulation), which was extended in a number of ways:

1. Simulating long sequences is disproportionately time-consuming because we must track many variants. We chose 0.5Mb as a good compromise between time efficiency and informativeness, and aimed for 500 replicate runs of each set of conditions, equivalent to about one tenth of the human genome. For a few key conditions such as the null model with constant mutation rate and no introgression, we extended to 1000 runs. Each run was initialized using a randomly selected segment from human chromosome 1 (build GR38, rejecting any sequences that contained ‘N’s), giving realistic initial base compositions and realistic frequencies for three-base motifs, crucially the XCG triplets. In all cases, we use a recombination rate of 2 × 10^−8^.
2. Our simulations begin with a single population that splits 250,000 generations ago (6 million years at generation length = 25) into a chimpanzee lineage, P1, and a lineage that will become Hominins, P2, both of size 50. We used small population sizes to reduce run times, and they are reasonable because this is effectively a burn-in period where we are interested only in substitutions, not diversity.
3. We added a Neanderthal population, P3, with size 5000 that splits from P2 about 20,000 generations ago, allowing us to add introgression where required. We also explored the impact of increasing the Neanderthal size to 10,000 (results from these runs are not included).
4. We used P2 effectively to seed the Gravel model, but with a reduced initial burn-in period of 19,000 generations, approximately four times the population size. Test runs with longer burn-in periods produced qualitatively indistinguishable results.
5. The Gravel model includes extremely low migration rates between all human population combinations. These were removed because our focus is on mutation-rate effects rather than gene flow and migration blurs mutation-rate asymmetries. Removing migration simplifies interpretation without altering the conclusions.
6. Basic mutation pattern. We used two mutation regimes: TRIPLTS, where every triplet has its own empirically determined mutation rate, and ONEMU, where we simulate only a difference between transitions and transversions. For maximum realism, in the TRIPLETS model we analysed the 1KG data and counted the numbers of each of the 64 possible triplets with a minor allele present in five or fewer copies across all samples, assumed to represent the products of recent mutations. Each count of polymorphic sites was then divided by the total number of sites with that triplet in the genome to obtain a measure of relative mutability. We then normalized each score so the mean null (unmodified) rate across all 64 triplets in the human genome matched the Gravel average mutation rate of 2.36 × 10^−8^. We specify ‘for the genome’ because overall mutation rate is a function of mutation rate x frequency across all triplets, so the absolute mutability of individual triplets means little unless coupled to their frequency.
7. In SLiM, mutation rate variation over time or across populations is achieved by discarding some proportion of mutation events according to user-specified criteria. Consequently, increases above the initialized mutation rate are not possible. To allow for a maximum 5X rate increase, we set the initialized rate to 5X the required null rate and, unless otherwise specified, discard 80% of mutations at random. This allows us to increase or decrease rates as desired within particular populations. Note, we do not attempt to model the complexities discovered by Harris and Pritchard (2017), where different triplet motifs have evolved at different rates across different populations.
8. Mutation rate variation. We modelled two forms of mutation rate variation. First, the mutation rate in humans either stays constant or increases linearly, beginning 5,919 generations before the present, coincident with the African expansion in Gravel et al.’s recipe. The terminal (present-day) human rates were set to 1X (no increase) or 2X the initial rate. Second, the mutation rate in non-Africans either stays the same as in Africa or drops to 0.25, 0.5, or 0.75 times the African rate.
9. In addition to changes in mutation rate, we explored introgression by adding a migration rate of 0 (no introgression), 0.001, 0.005, 0.01, or 0.015 from the Neanderthal lineage into the Eurasian lineage, 1974 generations, or about 50,000 years ago.
10. At the end of each simulation, we generated samples (one chimpanzee, one Neanderthal, and 50 each of Africans and Europeans, all diploid) and output base frequencies for each of the four taxa. Sites were retained if only two different variants were scored (an appreciable number of sites carried up to six alternate alleles, but as long as these were multiple instances of the same derived allele, they were retained). In this way, we mimic common practice among previous studies that focus only on biallelic sites. To allow study of mutational ancestry, we record the time and population of origin of all mutations.

Examples of the main and minor SliM scripts are given in ESM10, along with the simple bash scripts used to run them and generate the random initialisation sequences from human chromosome 1.

## Supporting information

ESM

## Data and script availability

All data and scripts needed to replicate our results and figures are available on GitHub (https://github.com/eelhaik/UnexpectedDtourAhead).

## Acknowledgment

EE was partially supported by the Swedish Research Council (2020-03485) and Erik Philip-Sörensen Foundation (G2020-011). The Swedish National Infrastructure for Computing (SNIC) at Lund provided the computing resources, partially funded by the Swedish Research Council through grant agreement no. 2018-05973. We are grateful to Mark Moore, Cecilia Padilla and Monty Slatkin for their constructive comments.

## Contributions

WA conceived the study. Both authors carried out the analyses and wrote the paper.

