## Supplementary material for "Unexpected D-tour Ahead: Why the D-Statistic, applied to Humans, Measures Mutation Rate Variation and Not Neanderthal Introgression": ESM

### ESM1. Transition:transversion ratios in relation to outgroup alleles

The number of sites that carry recurrent mutations can be assessed by considering transition-to-transversion (TS:TV) ratios. At polymorphic sites generated by a single mutation, the TS:TV ratio is expected to be around two across the whole genome (The International HapMap Consortium 2003; DePristo et al. 2011). However, because recurrent sites require two mutation events, the relative probabilities are multiplied, such that the expected TS:TV ratio is approximately squared, rising from two to around four. We exploit this theory to estimate the proportion of sites that contribute to D and carry recurrent mutations. For this, we estimated TS:TV ratios across the mutation spectrum (sensu Harris 2015; Harris and Pritchard 2017), using the central base in all possible three-base combinations = 'triplets'). To estimate which triplet mutated, we assume the human major allele represents the most likely human ancestral state. Under this assumption, we counted mutated triplets (i.e., triplets where the central base is polymorphic) across the genome, partitioned by the bases carried by four outgroup taxa: the Vindija and Altai Neanderthals, the Denisovan, and the chimpanzee (Table S1). Sample sizes were matched meticulously to try to avoid bias when determining the major allele (Africa = 168 alleles drawn from each of the five non-admixed populations, non-Africans = 56 alleles drawn from each of five populations from Europe, East Asia and South Asia). We note that some residual bias is unavoidable because differences in demographic history, population structure and relatedness influence allele-frequency distributions and hence the probability that the major allele represents the ancestral state.

Most outgroup conformations give TS:TV ratios a little over two. The glaring exception is the class of sites where all non-human Hominins carry the human reference allele which is also the human major allele and the chimpanzee carries the human alternate allele (orange column Table S1). Such sites likely result from one mutation between the chimpanzee and hominins plus a second mutation in humans, and the ratio reflects this, at approximately four. The probability of a recurrent mutation scales with the mutation rate squared, as reflected in the very large ratios seen at XCG triplets (highlighted in green), which include CpG motifs and have much higher mutation rates (Carlson et al. 2018). Here, ratios reach 22.3.

| taxa | R-A-R | A-A-R | R-R-A | A-R-A | RRRRR | ARRRR | RAAAA | AAAAA | RRRRA | ARRRA | RAAAR | AAAAR | RAARR | AAARR | RRRAA | ARRAA |
|---|---|---|---|---|---|---|---|---|---|---|---|---|---|---|---|---|
| minor | A | R | A | R | A | R | A | R | A | R | A | R | A | R | A | R |
| AAA | 1.27 | 1.29 | 1.38 | 1.03 | 1.27 | 1.24 | 1.27 | 1.24 | 1.41 | 1.24 | 1.26 | 1.19 | 1.26 | 1.14 | 1.33 | 1.10 |
| AAC | 2.09 | 1.15 | 1.09 | 1.93 | 2.24 | 1.08 | 1.09 | 2.08 | 1.31 | 2.19 | 2.36 | 1.13 | 2.28 | 1.03 | 1.09 | 2.09 |
| AAG | 1.63 | 1.58 | 2.08 | 1.89 | 1.71 | 2.20 | 2.34 | 1.77 | 2.28 | 2.08 | 1.86 | 2.32 | 1.74 | 2.29 | 2.10 | 1.99 |
| AAT | 3.18 | 2.13 | 2.54 | 3.13 | 3.44 | 1.91 | 1.96 | 3.37 | 4.04 | 3.17 | 3.52 | 2.51 | 3.41 | 2.09 | 1.93 | 3.63 |
| CAA | 1.67 | 2.50 | 2.52 | 1.62 | 1.47 | 2.98 | 2.80 | 1.45 | 2.82 | 1.86 | 1.56 | 3.09 | 1.44 | 2.91 | 2.71 | 1.44 |
| CAC | 1.90 | 3.79 | 2.80 | 2.40 | 1.67 | 3.19 | 2.97 | 1.78 | 3.24 | 2.38 | 2.00 | 3.55 | 1.80 | 2.73 | 2.98 | 1.73 |
| CAG | 2.13 | 4.94 | 4.70 | 2.33 | 1.98 | 5.01 | 5.33 | 2.05 | 5.45 | 2.69 | 2.54 | 6.02 | 2.07 | 4.49 | 5.04 | 2.42 |
| CAT | 2.86 | 4.55 | 5.84 | 2.65 | 2.77 | 6.19 | 5.35 | 2.74 | 7.13 | 3.47 | 3.01 | 6.23 | 2.79 | 6.17 | 5.53 | 2.76 |
| GAA | 1.41 | 1.84 | 1.72 | 1.30 | 1.36 | 1.66 | 1.76 | 1.45 | 1.80 | 1.43 | 1.45 | 1.62 | 1.38 | 1.84 | 1.53 | 1.40 |
| GAC | 1.80 | 2.14 | 1.73 | 1.84 | 1.52 | 1.83 | 1.67 | 1.57 | 1.74 | 1.88 | 1.71 | 1.75 | 1.80 | 1.46 | 1.55 | 1.40 |
| GAG | 1.43 | 2.28 | 3.25 | 1.13 | 1.37 | 3.09 | 3.24 | 1.39 | 2.96 | 1.50 | 1.58 | 3.01 | 1.47 | 2.28 | 3.02 | 1.39 |
| GAT | 1.29 | 2.15 | 2.02 | 1.44 | 1.37 | 2.21 | 2.08 | 1.44 | 2.46 | 1.62 | 1.49 | 2.19 | 1.31 | 1.81 | 2.18 | 1.62 |
| TAA | 1.49 | 0.92 | 1.10 | 1.75 | 1.78 | 1.78 | 1.69 | 1.70 | 1.52 | 1.61 | 1.63 | 1.39 | 1.69 | 1.55 | 1.67 | 1.87 |
| TAC | 3.00 | 1.62 | 1.82 | 3.39 | 3.02 | 1.88 | 1.71 | 3.12 | 2.31 | 3.23 | 3.05 | 1.84 | 2.88 | 1.67 | 1.84 | 3.02 |
| TAG | 2.31 | 2.73 | 3.81 | 2.42 | 2.51 | 4.24 | 4.10 | 2.44 | 4.76 | 2.52 | 2.66 | 4.24 | 2.59 | 4.60 | 4.12 | 2.40 |
| TAT | 3.11 | 1.94 | 2.22 | 3.79 | 4.20 | 2.49 | 2.22 | 4.44 | 3.58 | 4.28 | 4.15 | 2.82 | 4.26 | 2.32 | 2.32 | 4.49 |
| ACA | 1.73 | 2.06 | 1.62 | 2.03 | 1.95 | 2.05 | 1.84 | 2.00 | 2.15 | 1.96 | 2.01 | 1.91 | 1.86 | 2.49 | 1.84 | 1.99 |
| ACC | 1.28 | 1.57 | 1.50 | 1.27 | 1.37 | 1.94 | 1.82 | 1.26 | 1.86 | 1.13 | 1.15 | 1.93 | 1.18 | 1.92 | 1.78 | 1.08 |
| ACG | 12.55 | 8.10 | 11.86 | 13.34 | 14.20 | 3.73 | 4.16 | 11.63 | 22.31 | 11.90 | 11.88 | 9.01 | 12.27 | 5.38 | 5.43 | 9.52 |
| ACT | 1.69 | 2.81 | 2.61 | 1.71 | 1.64 | 2.74 | 2.67 | 1.51 | 2.91 | 1.47 | 1.63 | 2.83 | 1.50 | 2.78 | 2.78 | 1.56 |
| CCA | 1.66 | 1.23 | 1.58 | 1.43 | 1.94 | 1.71 | 1.51 | 1.83 | 1.93 | 1.97 | 2.05 | 1.58 | 1.94 | 1.53 | 1.61 | 1.80 |
| CCC | 1.88 | 1.71 | 1.61 | 2.26 | 1.76 | 1.70 | 1.52 | 1.62 | 2.16 | 1.87 | 1.90 | 1.84 | 1.73 | 1.62 | 1.68 | 1.77 |
| CCG | 9.82 | 4.18 | 7.95 | 10.41 | 10.85 | 3.13 | 3.49 | 9.21 | 17.14 | 8.56 | 10.04 | 8.07 | 10.59 | 4.08 | 4.29 | 10.49 |
| CCT | 1.95 | 1.60 | 1.57 | 1.84 | 2.02 | 1.59 | 1.67 | 1.99 | 2.07 | 2.14 | 2.08 | 1.81 | 2.13 | 1.85 | 1.73 | 2.13 |
| GCA | 1.26 | 1.31 | 1.43 | 1.05 | 1.44 | 1.41 | 1.27 | 1.38 | 1.63 | 1.51 | 1.44 | 1.52 | 1.41 | 1.16 | 1.29 | 1.26 |
| GCC | 1.52 | 2.28 | 1.74 | 1.17 | 1.57 | 1.38 | 1.42 | 1.54 | 1.80 | 1.45 | 1.49 | 1.64 | 1.49 | 1.32 | 1.58 | 1.38 |
| GCG | 9.12 | 3.49 | 5.35 | 5.72 | 8.65 | 1.88 | 1.95 | 7.88 | 11.04 | 6.21 | 7.86 | 5.02 | 8.30 | 2.30 | 2.79 | 9.35 |
| GCT | 2.11 | 2.13 | 2.43 | 1.52 | 2.05 | 1.84 | 1.96 | 2.06 | 2.30 | 1.75 | 2.02 | 2.03 | 1.95 | 2.18 | 2.01 | 1.97 |
| TCA | 1.40 | 1.25 | 1.38 | 0.99 | 1.46 | 1.30 | 1.31 | 1.37 | 1.50 | 1.25 | 1.44 | 1.31 | 1.44 | 1.32 | 1.30 | 1.23 |
| TCC | 1.59 | 1.28 | 2.32 | 1.78 | 1.71 | 1.80 | 1.92 | 1.46 | 2.42 | 1.24 | 1.46 | 2.18 | 1.43 | 2.12 | 2.31 | 1.39 |
| TCG | 8.96 | 3.02 | 7.47 | 7.33 | 9.61 | 2.36 | 2.33 | 8.58 | 11.05 | 7.76 | 8.11 | 4.58 | 8.75 | 2.73 | 2.96 | 7.64 |
| TCT | 0.96 | 1.40 | 1.67 | 0.91 | 0.93 | 1.39 | 1.55 | 0.76 | 1.48 | 0.72 | 0.84 | 1.30 | 0.83 | 1.32 | 1.67 | 0.72 |
| nTV | 20 | 2 | 11 | 3 | 3,339 | 21 | 80 | 149 | 106 | 12 | 61 | 14 | 74 | 5 | 25 | 8 |
| nTS | 44 | 4 | 30 | 6 | 7,465 | 47 | 172 | 317 | 423 | 26 | 137 | 38 | 162 | 11 | 58 | 17 |
| **ALL** | **2.18** | **2.18** | **2.68** | **2.07** | **2.24** | **2.18** | **2.16** | **2.12** | **3.98** | **2.21** | **2.24** | **2.67** | **2.18** | **2.24** | **2.28** | **2.12** |
| % sites | 0.2 | 0.0 | 0.1 | 0.0 | 25.9 | 0.2 | 0.6 | 1.2 | 0.8 | 0.1 | 0.5 | 0.1 | 0.6 | 0.0 | 0.2 | 0.1 |
| % sites | 0.3 | 0.0 | 0.2 | 0.0 | 57.9 | 0.4 | 1.3 | 2.5 | 3.3 | 0.2 | 1.1 | 0.3 | 1.3 | 0.1 | 0.4 | 0.1 |

Table S1.1. **Transition-to-transversion ratios for triplets in relation to alleles carried by the outgroups**. *Taxa are ordered: human, Vindija, Altai, Denisovan, chimpanzee in red. R = human reference allele, A = human alternate allele, '-' = any. Minor is the minor allele in Africans and non-Africans or, where these differ, the minor allele in the pooled sample. XCG transitions (green), sites most likely to carry recurrent mutations (chimpanzee = the human minor allele = the alternate allele) (orange), intersection (pink) and overall ratios (blue) are emphasised. Total numbers of sites (nTV, nTS) are given in 1000s. The bottom rows give the percentage of all sites.*

We note that the widely cited TS:TV ratio of two is a genome-wide average (DePristo et al. 2011) that conceals substantial variation in substitution rates among sites and sequence contexts (Aggarwala and Voight 2016). Even setting aside recurrent mutations, the different triplets vary greatly, with many ratios barely above one and, in some cases (e.g., AAA), ratios across all outgroup scenarios that do not exceed 1.5.

In addition to counting transitions and transversions, we also counted the number of D-informative sites per non-overlapping 50Kb window (Fig. S1.1). Of course, the actual number in each window depends on the populations being analysed and the sample sizes of alleles drawn from each population, with larger sample sizes and less related populations both tending to reveal more sites where at least one population is polymorphic. We also counted sites that are polymorphic in humans and where the Altai Neanderthal allele differs from the chimpanzee, and the results are nearly identical (the main hump shifts 1.5 units to the left).

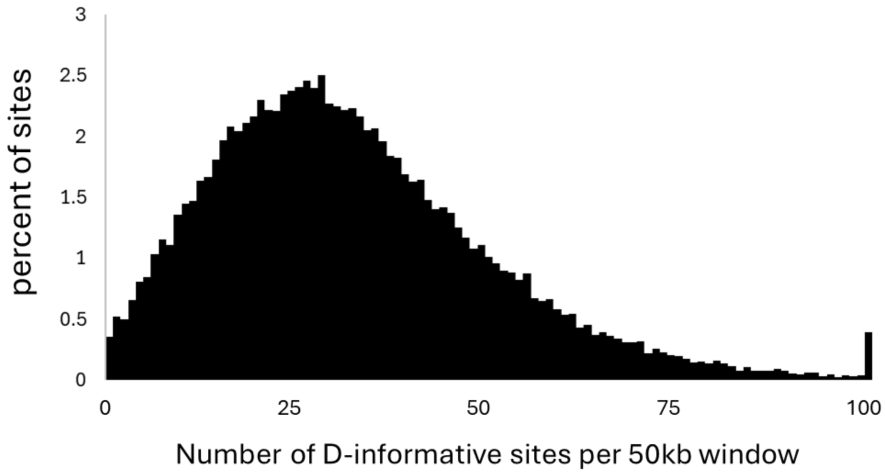

**Figure S1.1. Number of D-informative sites in non-overlapping 50kb windows.** *D-informative sites are defined by the qualifying outgroups: we counted all sites where the Altai Neanderthal and chimpanzee differ. For any given population pair, the number of sites will be somewhat lower but not massively so.*

### ESM2. Explaining variation in D across triplets

We have shown that D varies dramatically across the mutation spectrum, differing not only by triplet identity but also, profoundly, by whether the mutation is a transition (TS) or a transversion (TV) (Fig. 2, main text). One possible influence is the broader sequence context. One of the most important sources of context variation is likely to be CpG islands (Carlson et al. 2018), regions with elevated GC content and more CpG motifs than expected by chance. To test whether such regions affect D, we re-analysed the data, this time classifying sites as lying inside or outside CpG islands. Remarkably, although this classification causes no appreciable change to overall D, it massively increases the level of variation between triplets. Moreover, when we serendipitously compared the result of a trial run based only on chromosome 1 with the overall pattern, the plots differed greatly, with the single-chromosome plot showing peaks and troughs with far greater amplitude, many of which did not match their equivalents in the overall plot (Fig. S2.1).

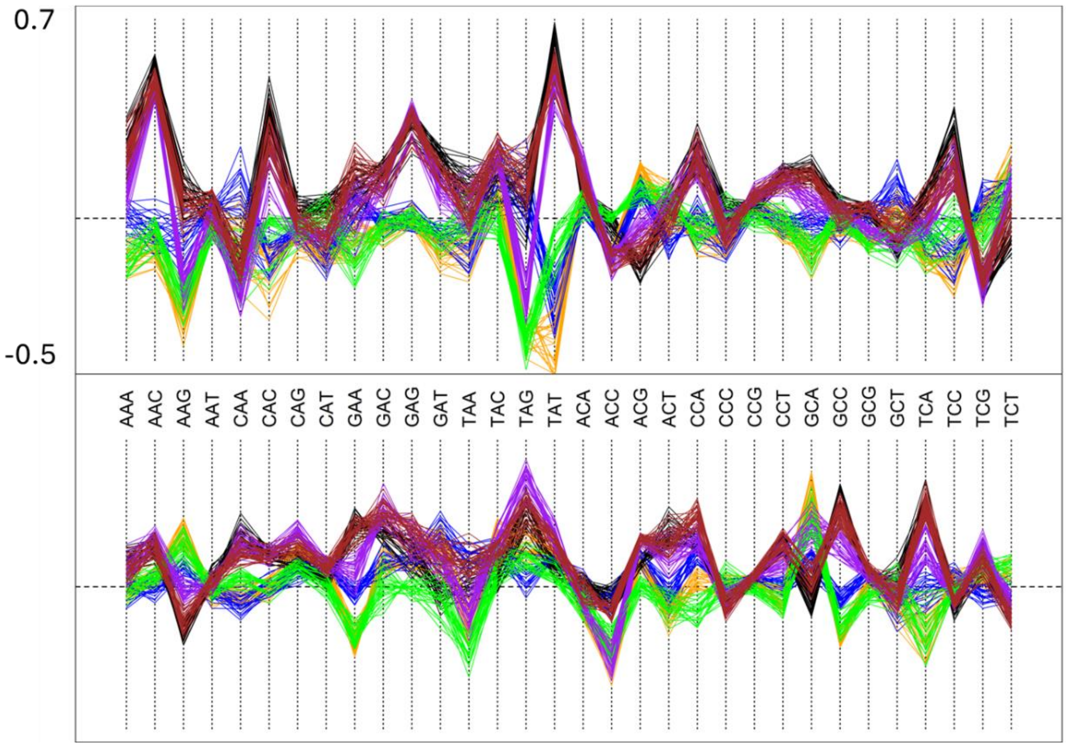

**Figure S2.1. Variation in D across the mutation spectrum in CpG islands.** *Data are for chromosome 1 alone. Triplets were classified according to the inferred ancestral triplet, the human minor allele assumed to be the derived state, and the human major allele assumed to be ancestral. We calculated D separately for each triplet, partitioned into* **TV** *(top panel) and* **TS** *(bottom panel). All possible pairwise comparisons are made between African (AFR), European (EUR), East Asian (EAS) and South Asian (SAS) populations: black (EUR-AFR); brown (SAS-AFR); purple (EAS-AFR); green (SAS-EAS); orange (EUR-EAS); blue (EUR-SAS). Every line is a different population comparison. D on the y-axis is the same in both panels and ranges from -0.5 to 0.7. This figure should be compared with Fig. 2 in the main text, where D values cover a range that is more than 8.5X smaller (-0.02 to 0.12).*

To learn more about this puzzling triplet behaviour, we generated individual chromosome plots of triplet against D for the largest nine chromosomes (Fig. S2.2). These reveal a completely unexpected pattern: each chromosome appears to have its own signature pattern. On the one hand, this provides a proximate explanation for why D is smaller and less variable across the mutation spectrum when chromosomes are combined: every time a triplet gives high D on one chromosome and low D on a second, the peak and trough cancel each other out. On the other hand, there is no obvious explanation for why chromosomes should differ so profoundly. It remains unclear why the profiles differ so greatly between chromosomes, or how D can reach such extreme values. However, these questions lie beyond the scope of this manuscript and must be the subject of future work.

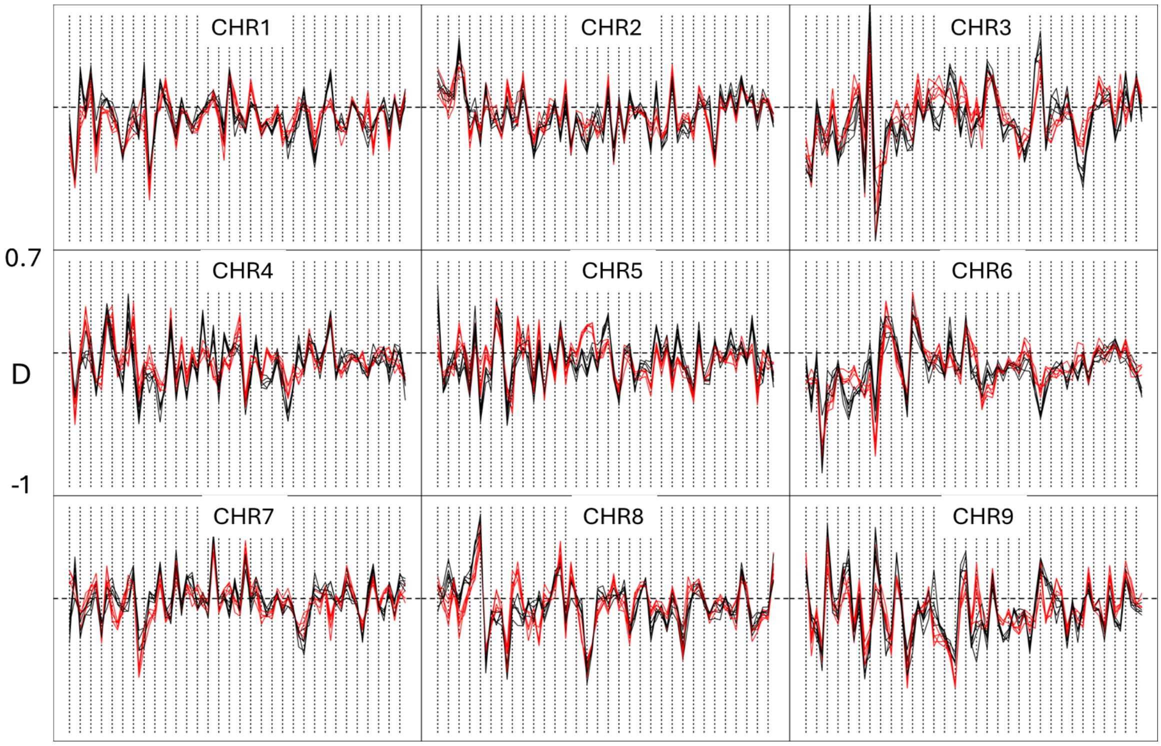

**Figure S2.2. Variation in D by triplet for the nine largest chromosomes.** *Each panel presents data for how D varies by triplet (32 TV followed by 32 TS in the usual order) at sites within CpG islands. Red and black lines are for five independent Europe-Africa and five independent East Asia-Africa population comparisons. By independent, we mean that no population is used more than once in each block of five. The y-axis scale is the same for all panels, and vertical lines are drawn for every second triplet to aid comparative interpretation.*

### ESM3. Relationship between triplet identity, flanking heterozygosity and D

Some triplets become more mutable and others less mutable with increasing flanking heterozygosity (Amos 2019). If changes in mutation rate drive D, then D will also be closely related to local heterozygosity. Consequently, we extended the prior study (Amos 2019), this time calculating mean heterozygosity within 1kb of each D-informative site and expressing it as the proportion of triplets in each heterozygosity bin that each triplet represents. In other words, we determined the extent to which each triplet tends to be either over- or under-represented relative to the flanking heterozygosity. We partitioned data by whether the site did (red) or did not (black) lie in a CpG island (Fig. S3.1). We confirm that regions that differ in heterozygosity also differ in triplet proportions. Moreover, for any given triplet, the trends can differ markedly between otherwise equivalent regions inside and outside CpG islands. Some of the strongest trends are seen for transitions at triplets beginning with ‘CA’ (outlined in green), and particularly for CAT where the two trends are diametrically opposing. The more common instances are those in which individual triplets are enriched within islands than outside of them, or vice versa, as indicated by the large separation between the red and black distributions (e.g. AAT transitions, panel 5 down and 4 across; AAA transversions, top-left panel).

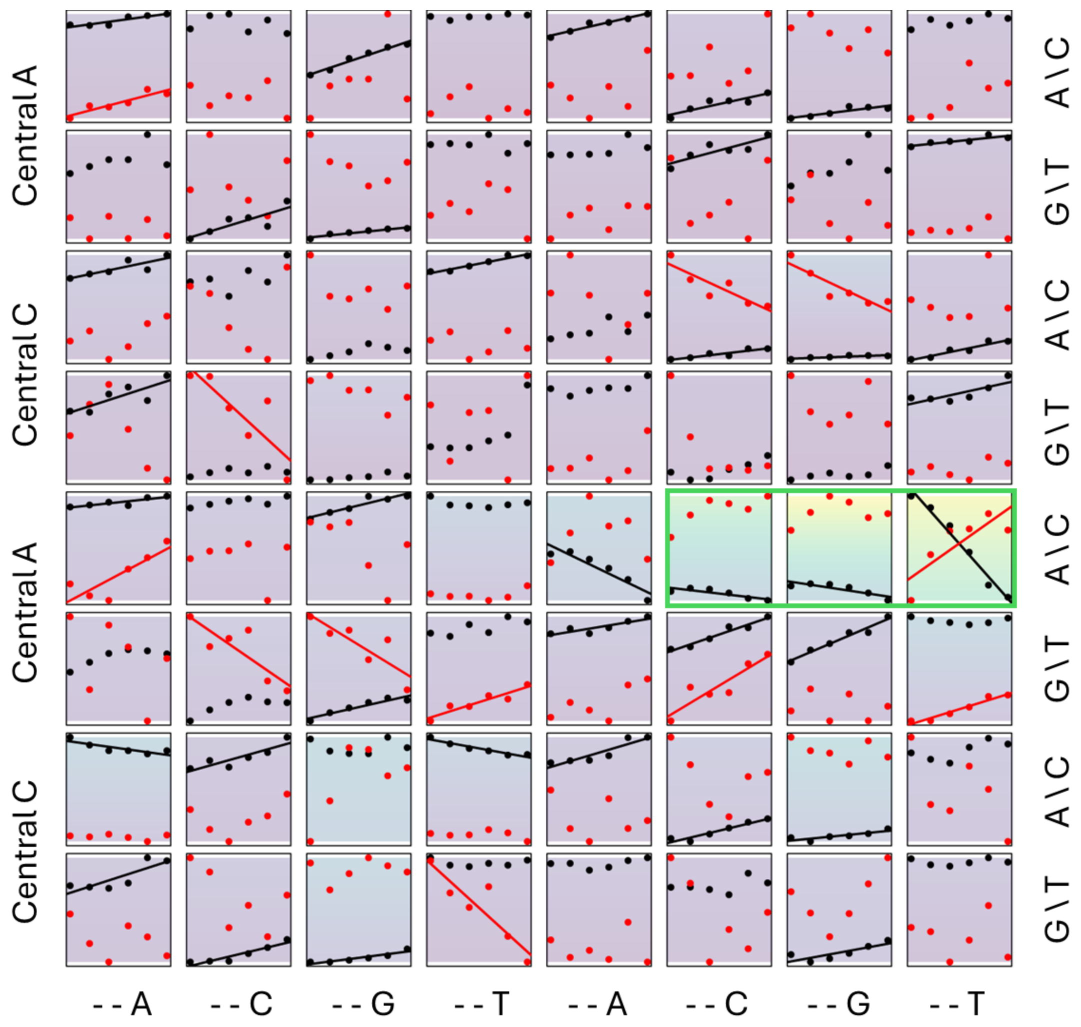

**Figure S3.1. Dependence of the mutation spectrum on local heterozygosity.** *The x-axis is six evenly spaced heterozygosity bins (methods). The y-axis is the proportion of all triplets of a given type found within a given heterozygosity bin, adjusted separately for each panel. Where trends are significant at p<0.05 (linear regression of the mean values, N=6), a trendline is included. For clarity, y-values have been replaced with a colour scale that ranges from zero (light purple to blue, green to yellow, with the maximum at 0.1. Thus, the highest representations are outlined in green and are for transitions at CCC, CCG and CCT. Triplets are coded by the 5’ base (two blocks of four in each row, listed to the right), the 3’ base (listed underneath), and the central base (indicated to the left). Thus, the top row of panels represents triplets: AAA, AAC, AAG, AAT,*

*CAA, CAC, CAG, CAT. The top four rows depict* TVs *and the bottom four* TSs. *Data for sites outside CpG islands are in black and inside CpG islands in red. Across 128 data series, 47 give significant regressions at P<0.05. Transversions at three triplets with initial 'CA' give unusually high values and exhibit large differences between CpG and non-CpG sites and are outlined in green.*

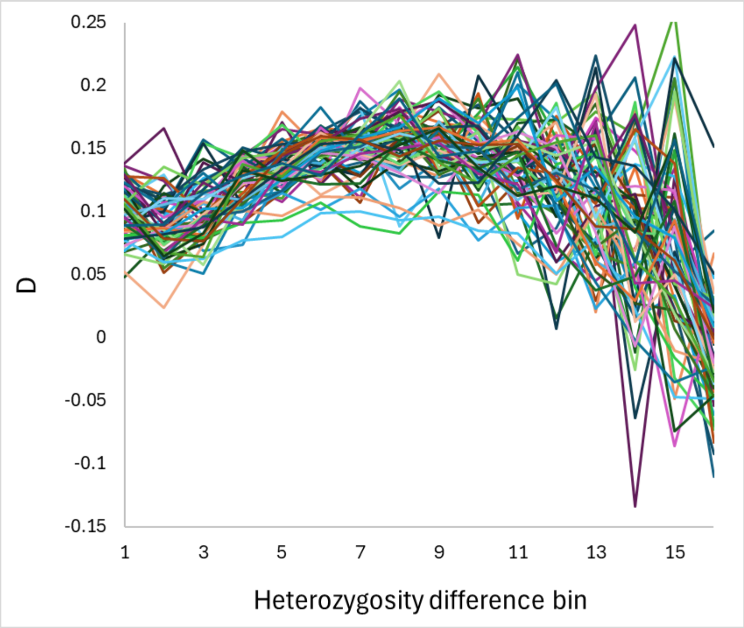

**Figure S3.2. Dependence of D on heterozygosity difference between Africa and Europe, calculated separately for each triplet.** *This plot shows the raw data used to compare similarity among the 64 profiles. Far fewer sites lie within CpG islands, but within the far greater scatter, an essentially identical pattern is seen.*

Because the probabilities of each mutation type often show strong and variable dependence on sequence context in terms of both GC content and local heterozygosity (Figs. S3.1, S3.2), it is also worth exploring whether D varies with heterozygosity. Therefore, we calculated D for each triplet and plotted it against the Africa–Europe heterozygosity difference. Even though different triplets on average give different D, in general all triplets give similarly shaped profiles with D increasing with heterozygosity difference up to a point and then decreasing thereafter (Fig. S3.2). To view these trends more clearly, we next plotted the average profiles, superimposing the sample size (ABBA + BABA) for each heterozygosity difference bin (Fig. S3.3).

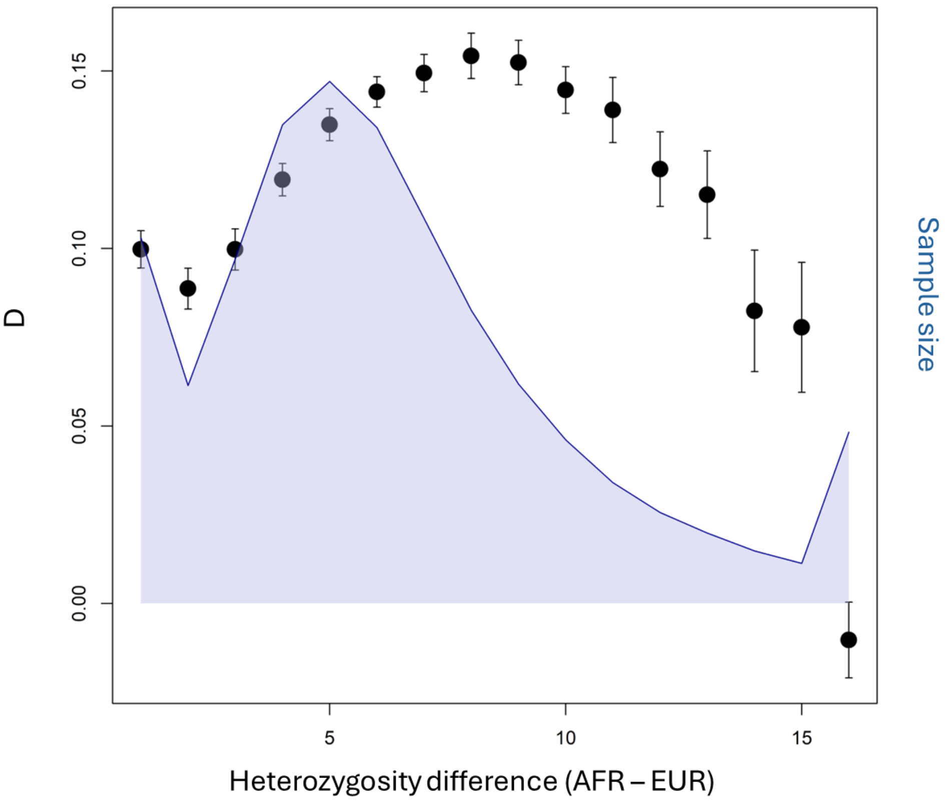

**Figure S3.3. Variation in D with heterozygosity difference between Africa and Europe.** *Error bars are +/- 2 standard errors of the mean, based on the 64 values for the different triplets. The relative sample size (area plot) is also shown.* Although the decline in D as heterozygosity difference increases beyond around eight appears convincing, across most sites D increases with *heterozygosity difference.*

To better understand why the relationship between heterozygosity difference and D is humped, we revisited the idea depicted in Fig. 1 (main text), where we point out how recurrent mutations that are more likely in Africa can drive both positive and negative D. On the common BBBBA background where the chimpanzee differs from all other taxa, recurrent B - > A mutations in Africa create ‘ABBA’s and drive positive D, suggesting introgression outside Africa. However, on the ABBBA background, where humans and chimpanzees carry the ancestral A allele and all non-human Hominins carry a derived variant, recurrent A -> B mutations in Africa create ‘BABA’s and hence drive a negative D that would imply introgression into Africa. This is exactly what we find. Partitioning the heterozygosity-D data by the alleles carried by the outgroups, we find diametrically opposing trends. When all three non-human Hominins carry the same allele (red/orange symbols in the left-hand panel), D increases more or less linearly with heterozygosity difference, and is largest where Africans have much higher heterozygosity than non-Africans. However, at sites where the hominins are polymorphic for either PXBXA (blue symbols) or PBBAA (black/grey symbols), the relationship goes in the opposite direction (Fig. S3.4). A changing balance between these two trends provides a proximate explanation for the humped trend seen in Fig. S3.3. At the same time, we find a strong trend in which relative mutation rate declines as African excess heterozygosity increases (right-hand panel).

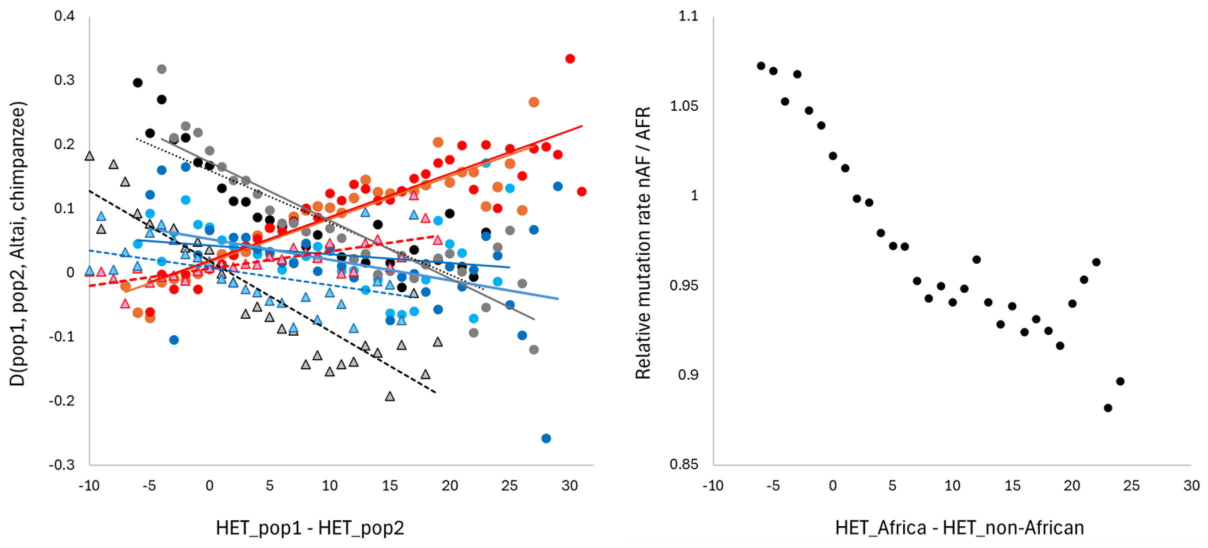

**Figure S3.4. Relationship between D, heterozygosity and the outgroup sequences.** *The 1KG data aligned to the Vindija, Altai, Denisovan and chimpanzee were analysed in non-overlapping 50kb windows. We chose three representative regions, taking 100 chromosomes from each of five populations from Africa (LWK, ESN, YRI, GWD, MDE), Europe (GBR, FIN, CEU, IBS, TSI) and East Asia (CHB, CHD, CDX, KDV, JPN). In each window, we calculated heterozygosity in each region, D(pop1, pop2, Altai, chimpanzee) and relative mutation rate.* ***Left***: *For D, each pairwise comparison (AFR-EUR; AFR-EAS, EUR-EAS, represented as darker colour, lighter colour and triangles, respectively) was calculated such that positive values would classically be interpreted as introgression into pop2. D-informative sites were divided into sites where all three non-human hominins have the same allele (red / orange); sites where the two Neanderthals differ from both the Denisovan and the chimpanzee (black / grey); and all remaining sites where the Altai and chimpanzee differ (blue).* ***Right***: *We also calculated relative mutation rate, using the Altai Neanderthal as the assumed human ancestral allele. For relative mutation rate, we used samples of 168 chromosomes from each of the five African populations and 56 chromosomes from each of the 15 non-African populations in Europe, East Asia and South Asia, giving the same number for each continent. The x-axis shows heterozygosity differences in integer bins, calculated as the sum of heterozygosity differences between pop1 and pop2 over all informative sites. For clarity, we excluded the highly variable 2.5% of data at each end of each trend.*

The above trends appear consistent with the model we suggest, with PBBBA sites enriched for ancestral B in humans and PBBAA/PXBXA sites enriched for ancestral A in humans. By contrast, the opposing trends in D appear impossible to reconcile with a model based on introgression. Across the entire genome, the contribution of introgressed fragments to heterozygosity would be tiny relative to the large changes driven by the out-of-Africa bottleneck. Across the genome, any given window that carries an introgressed fragment could have much increased heterozygosity. However, as we show (Fig. 8, main text), the resulting trend affects only the small minority of windows with higher heterozygosity outside Africa. Consequently, an introgression-based explanation does not fit the data: it would predict a single trend rather than two opposing trends, and the main trend it would produce runs opposite to the main trend observed empirically. Under introgression, high D should occur where non-African heterozygosity is high because regions carrying more introgressed Neanderthal variation should contain both more archaic-derived variation and greater excess allele sharing (Green et al. 2010), but empirical data show high D where excess heterozygosity in Africa is greatest.

We note also that the striking difference between PBBBA and PXBXA sites is seen despite a sample size of only two Neanderthals and one Denisovan. Clearly, larger sample sizes of archaics will act to move some proportion of the PBBBA sites into the PXBXA category, while more Denisovans, in particular, would help identify sites carrying Neanderthal-specific mutations, i.e., the PBBAA sites (black / grey). We note that even the difference between PBBAA and PXBXA sites is consistent with our model. At PBBAA sites, the most likely scenario is an ancestral A allele and a derived B in Neanderthals, matching the middle row in Fig. 1 (main text), hence the strong negative trend. At PXBXA sites (blue), there is far more ambiguity about the ancestral state, so the opposing signals are more mixed and the trends much weaker overall. Finally, as would be expected, entirely non-African comparisons (triangles) generally show smaller magnitude D values and weaker trends (Fig. S3.4).

As mentioned, the right-hand panel of Fig. S3.4 shows a strong trend toward the non-African mutation rate becoming progressively lower relative to the African rate as excess heterozygosity in Africa increases. This replicates the finding in Amos (2013). However, the earlier study used a measure of relative mutation rate based only on rare alleles, which likely makes it less robust to demographic differences. In the current analysis, we use a measure based on all sites where all outgroups have been called (see ESM4 below). Note how the relative mutation rate switches from higher outside Africa to higher inside Africa as the heterozygosity difference switches from positive to negative: although very few windows show higher heterozygosity outside Africa, where they do occur, mutation rate outside Africa is also higher. These results contrast with simulated data, where there is no discernible link between heterozygosity difference and relative mutation rate (Fig. 8, main text)

### ESM4. Ancestral state assignment in relative mutation rate estimation

Previous attempts to estimate the relative mutation rate in one human lineage compared with another have assumed that the chimpanzee is an adequate surrogate for the true human ancestral state (e.g., Harris and Pritchard 2017). In a three-way alignment of human1, human2, and chimpanzee, we count sites of form BAA and ABA as derived alleles in human1 and human2, respectively. However, any mutations on the long branch separating chimpanzees from humans will reverse the assumed ancestral and derived states in humans, generating an inverted signal (Fig. S4.1). In addition, human data are strongly influenced by an observation bias created by assigning one allele, typically the commoner one, as the reference allele, which is also more likely to be the human major allele and therefore the human ancestral allele (ESM 8).

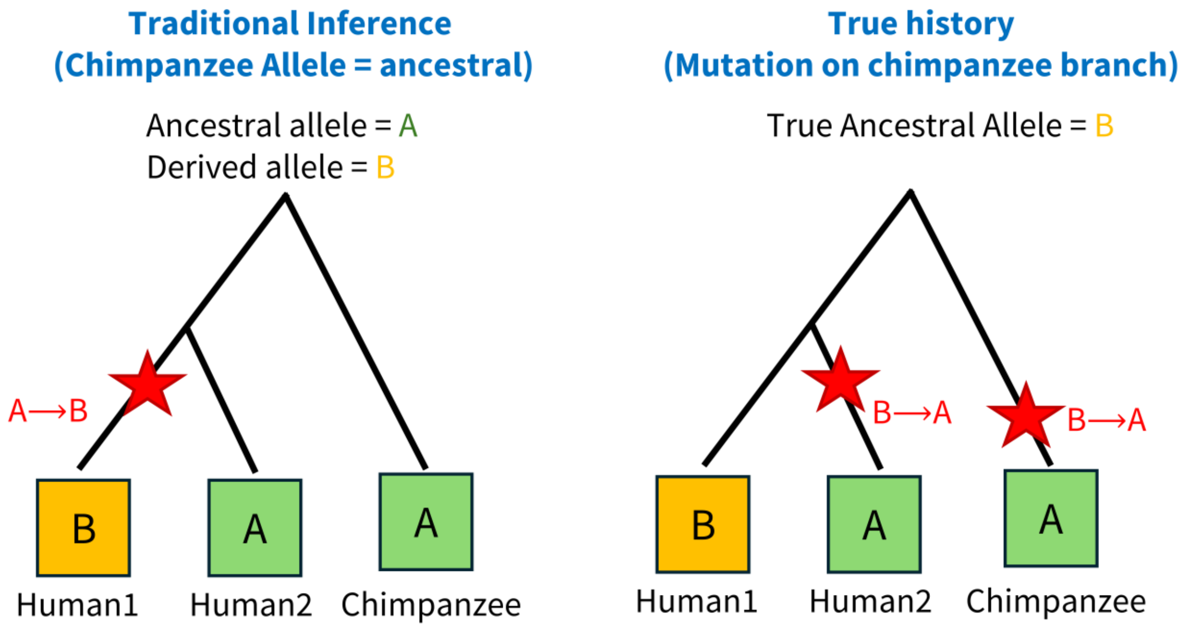

**Figure S4.1. Chimpanzee polarization can reverse the inferred direction of mutation.** *The observed alignment, BAA, is identical in both panels. Under the traditional interpretation (left), the chimpanzee allele is correctly assumed to represent the ancestral human state, and a mutation event is credited to the human1 lineage. However, when a mutation occurs between chimpanzees and humans (right), the assumed ancestral state is incorrect, causing mutations to be credited to the wrong branch. Thus, recurrent mutations can cause mispolarisation.*

To mitigate the impact of the long branch to the chimpanzee, we explored using non-human Hominins as the outgroup. Specifically, we required all three individuals (Vindija, Altai, Denisovan) to carry the same allele. In simulated data, without introgression and based on the default Gravel model with the full matrix of triplet mutation rates, this adapted method works well (Table S4.1). Where there is no mutation rate difference between Africans and non-Africans, the relative rate is effectively one. When the non-African mutation rate is reduced, the ratio falls progressively in proportion,

| Drop | Rise | Triplets | OneMu |
|---|---|---|---|
| **0** | **1** | 0.998 | 1.000 |
| **0** | **2** | 0.990 | 0.995 |
| **0.25** | **1** | 0.960 | 0.964 |
| **0.25** | **2** | 0.922 | 0.927 |
| **0.5** | **1** | 0.923 | 0.930 |
| **0.5** | **2** | 0.850 | 0.850 |
| **0.75** | **1** | 0.881 | 0.884 |
| **0.75** | **2** | 0.779 | 0.777 |

**Table S4.1. Relative mutation rate estimation in simulated data.** *Data are for 500 replicate runs each of a 500Kb segment using the Gravel model. We compare versions where each triplet has its own mutation rate (Triplets) with a simple two-rate model in which transitions are twice as likely as transversions. Relative mutation rate is estimated assuming the Neanderthal allele is the human ancestral state. Drop is the proportional reduction in the non-African rate relative to Africans (0.25 = 25% lower), and Rise determines whether human mutation rates are constant (Rise=1) or increase linearly by a factor of two over the last 6,000 generations (Rise=2).*

We next applied this adapted method to empirical data. Overall, the results suggest that the non-African mutation rate is approximately 10–20% lower than the African rate, although individual triplets vary significantly and consistently around the mean (Fig. S4.3), with values ranging from under 0.9 up to around 1.15. We confirmed that this variation reflects genuine variation in the underlying relative mutation rates both through the high levels of consistency between independent pairwise population comparisons (Fig. S4.3) and because the ratios for representative but independent Africa – Europe and Africa – East Asia comparisons are correlated regardless of whether the chimpanzee state is ignored or required to match the non-human Hominins (Fig. S4.4).

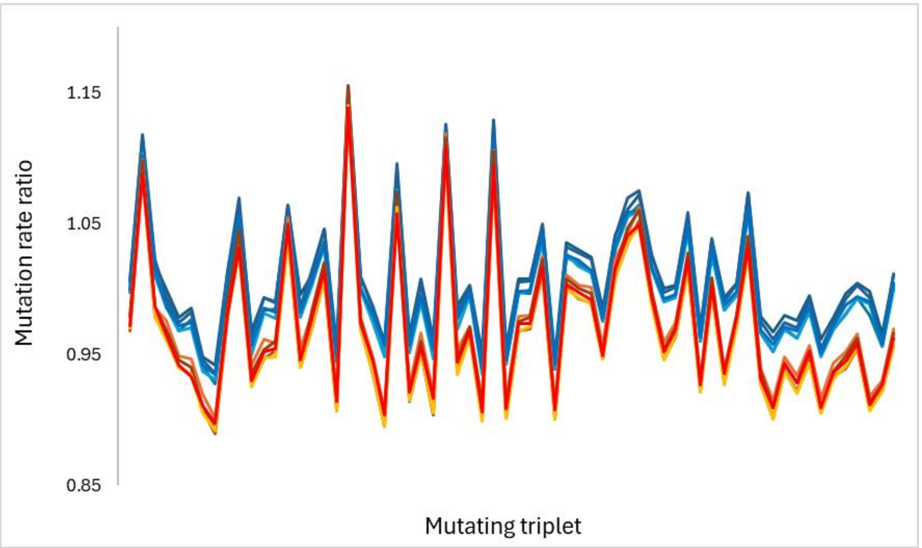

**Figure S4.3. Variation in mutation rate ratio across the mutation spectrum.** *The mutation rate ratio was calculated separately for every triplet (32 TV then 32 TS, triplet order as elsewhere) for five independent pairwise Europe – Africa comparisons (GBR-LWK, FIN-ESN, CEU-MSL, IBS-GWD, TSI-YRI, blue lines) and for five independent East Asia – Africa comparisons (CHB-LWK, CHS-ESN, CDX-MSL, KHV-GWD, JPT-YRI, red/orange lines). By independent, we mean that no population occurs more than once in each block of five. We calculated the relative mutation rate using the method of Mallick et al. (2016) , but with the assumed human ancestral state based on the three non-human Hominins rather than the chimpanzee.*

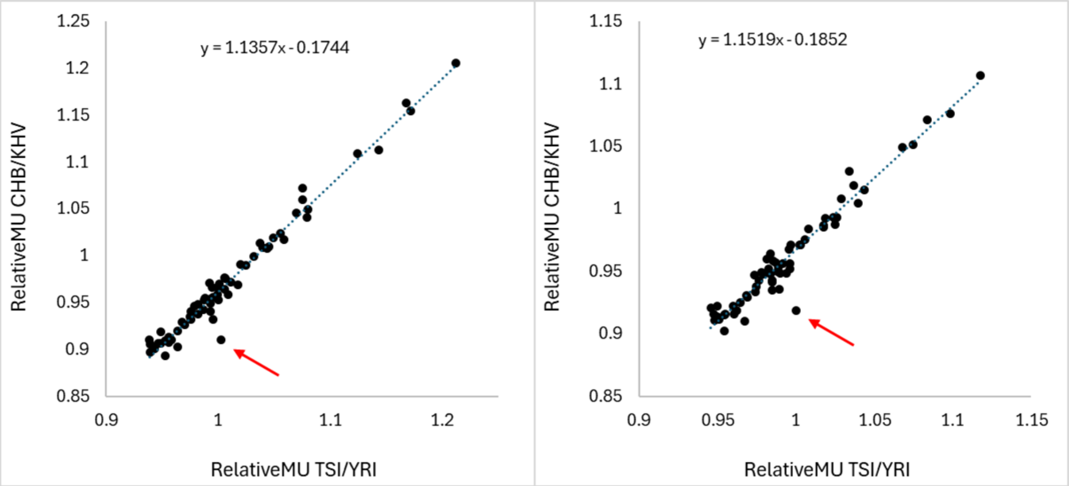

**Figure S4.4. Correlation between estimates of relative mutation rate across the mutation spectrum.** *We compare two maximally dissimilar African–non-African population pairs: East Asia (CHB-LWK) and Europe (TSI-YRI). Relative mutation rates were calculated assuming the non-human Hominins carry the human ancestral allele. The left-hand plot uses only sites where all non-human taxa carry the same allele. The right-hand panel uses all sites where the Altai, Vindija and Denisovan carry the same allele, regardless of the chimpanzee allele. In both cases, transitions at triplet TCC, which Harris and Pritchard (2017) identified as having an unusually high mutation rate in Europeans, produce a large outlier (red arrow).*

We next applied this method to test whether heterozygosity differences between populations predict corresponding differences in mutation rate. Here, we needed to ensure complete independence between relative mutation rate and D. Consequently, when estimating relative mutation rate, we required all outgroups to carry the same allele, including the chimpanzee. Sites that contribute to D always differ between Altai and chimpanzee, while sites that contribute to relative mutation rate always match

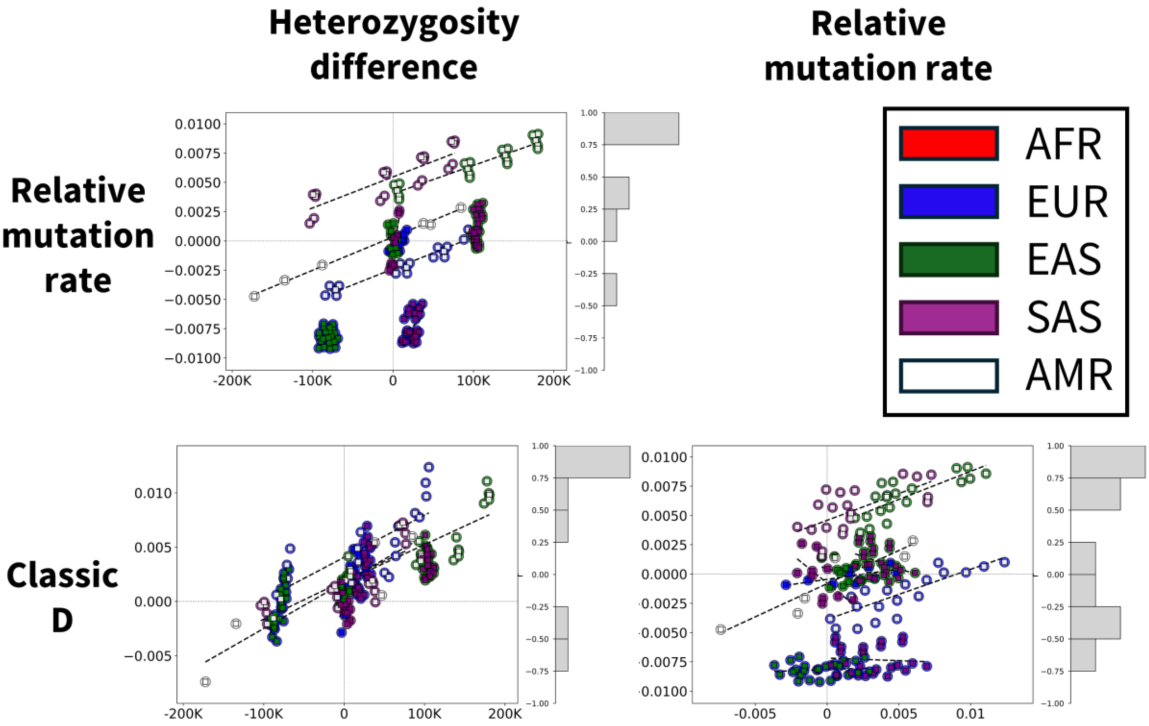

**Figure S4.5**. *Relationships among heterozygosity difference, the D statistic, and relative mutation rate among non-Africans. Pairwise scatterplots compare heterozygosity difference, the classic D statistic, and the mutation-rate statistic across 1KG populations. Each point represents a population pair and is coloured by geographic region. Most regressions (dashed lines) are positive (see histograms for r distribution). Although relative mutation rate is usually expressed as MU_Eur / MU_Afr, for these plots we used a statistic that is directly analogous to D: (MU_Pop1 – MU_Pop2) / (MU_Pop1 – MU_Pop2), where Pop1 = Africa / the population closest to Africa. Positive values therefore indicate a higher rate in Africa. In these plots, we use only sites where the archaics all carry the human reference allele. Hisotgrams alongside each plot give the distributions of r values.*

between Altai and chimpanzee; the sites used to calculate the two measures are completely non-overlapping. When plotted against heterozygosity difference, we find ubiquitous and highly significant relationships both overall (Fig. 7, main text) and within each major geographic region (Figs. S4.5, S4.6). We found that heterozygosity difference, relative mutation rate, and the D statistic are all strongly correlated. Pairwise comparisons with larger heterozygosity differences exhibit larger mutation-rate differences, and both quantities predict D. Together, these relationships support a common underlying process in which the reduction in heterozygosity following the Out-of-Africa bottleneck caused a parallel reduction in mutation rate, which in turn generated the observed D signal.

Of course, we can never know the true ancestral allele with certainty. However, replacing the chimpanzee with an archaic consensus offers a logical and effective improvement that considerably reduces the impact of recurrent mutations, a problem previously overlooked. Nonetheless, the method remains imperfect because recurrent mutations persist, and wherever they occur, they tend to generate an inverted signal. This problem is exacerbated by the truism that the higher a site's mutation rate, the more likely it is to contribute to mutation-rate estimation, but also the more prone it is to recurrent mutations.

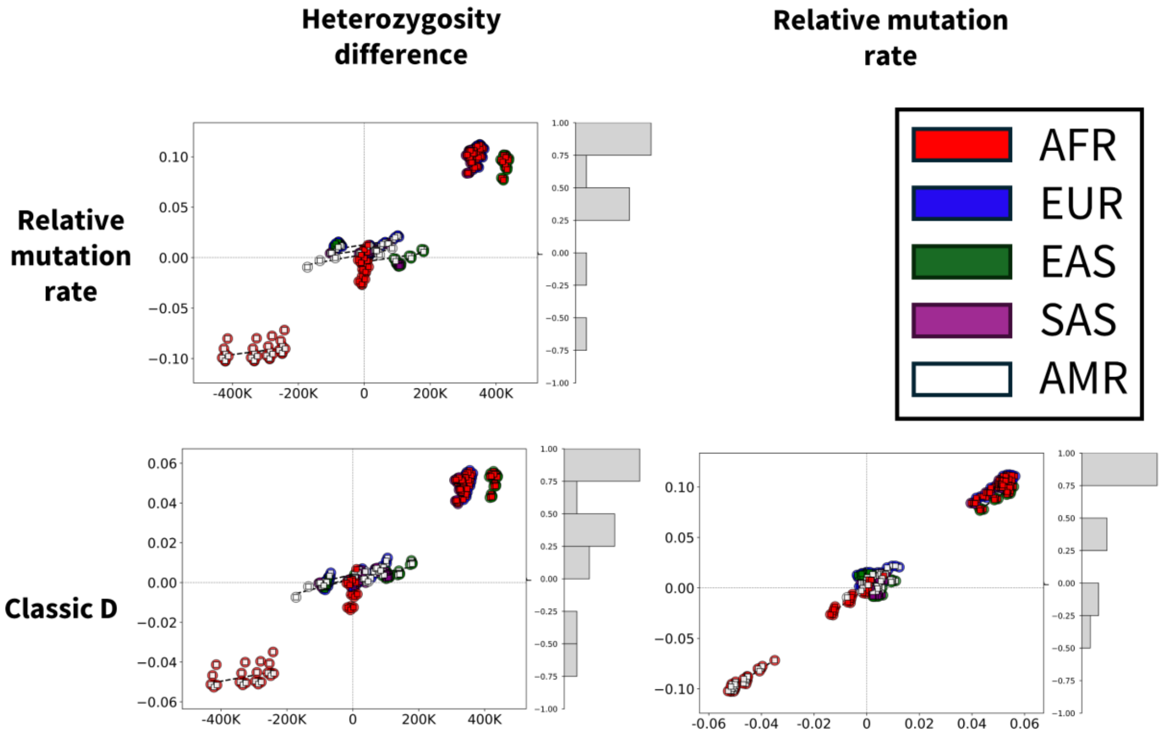

**Figure S4.6. Relationships among heterozygosity difference, the D statistic, and relative mutation rate among the major regions.** *Pairwise scatterplots compare heterozygosity difference, the classic D statistic, and relative mutation rate statistic across 1KG populations. Most regressions (dashed lines) are positive (see histograms for r distribution). Relative mutation rate is calculated as an analogue of D: (nP1 – nP2) / (nP1 + nP2), where nP1 and nP2 are the numbers of derived variants assigned to populations 1 and 2 and it uses only those sites where all non-humans carry the human reference allele. Each point represents a population pair and is coloured by geographic region. Dashed lines indicate least-squares regression fits. Hisotgrams alongside each plot give the distributions of r values.*

Note how, compared with the same plots based on all sites (Fig. 7, main text), all regressions are much tighter, particularly for D vs relative mutation rate. This alone creates what appears to be an unanswerable challenge to the introgression hypothesis. Relative mutation rate cannot have anything to do with introgression because all non-humans carry the same allele and it is calculated from an entirely non-overlapping sub-set of sites. What possible mechanism, then, could drive a correlation with D? We are unable to think of any. Although correlation does not imply causation, neither does it imply the absence of causation. Where a robust correlation exists, the alternatives include direct causation, reverse causation, a common cause or coincidence. If the latter alternatives provide no plausible explanation for the observed association, a causal relationship becomes the parsimonious interpretation. Moreover, correlations are not seen in simulated data with introgression and constant mutation rates. In contrast, a strong correlation is predicted by and indeed an unavoidable consequence of the MRVH. Here, the two measures are calculated in exactly the same way and differ only in the conditioning on which alleles are carried by the outgroup taxa: if all outgroups are the same, the measure is relative mutation rate, while if the chimpanzee and Altai differ, we get D. Consequently, wherever mutation rate varies between populations, D will also covary. However, if mutation rate does not vary between populations, relative mutation rate will always be one and the two measures cannot covry. Remarkably, Green et al. (2010) recognised how similar are these two measures, but dismissed the evidence for a mutation rate difference as being due to sequencing errors, despite being based on 10X the number of sites used to calculate D (p139 in their ESM)!

Finally, sites where all non-humans carry the human reference allele account for 83.8% of all sites (Table S1.1 above), and these sites indicate that the mutation rate outside Africa is approximately 80% of the rate seen in Africans. Since the relative rate calculated over all sites is around 93-95%, the 13% of sites where all non-humans carry the alternate allele must generate a strong counter-signal that cancels out much of the first signal. It is tempting to invert the counter-signal and add it to the 80%, giving a relative rate well below 80%. However, this is not valid because some of the apparent rate drop likely reflects a combination of differences in allele frequency spectra between Africans and non-Africans, coupled with biased sub-setting of sites. Nonetheless, we feel that a relative rate of around 80% is unlikely to be far from the true value.

### ESM5. Decomposing D according to which allele matches the human major allele

Setting aside the likely rare instances of DNA sequences under strong positive selection, archaic DNA fragments entering humans at low levels through interbreeding around 50,000 years ago will almost all have remained below 20% in frequency (Green et al. 2010). Introgressed alleles will therefore almost invariably match the human minor allele, and D calculated using only sites where the Neanderthal allele matches the human minor allele should be larger than D calculated over all sites. We tested this prediction by partitioning D according to whether the human major allele matches the chimpanzee or the Neanderthal. We also explored the impact of using only sites where the three archaics (Vindija, Altai, Denisovan) all carry the same allele, implying that this allele is ancestral, or are polymorphic, implying the presence of a derived variant created by a mutation that occurred in hominins but outside humans. As usual, we also partitioned the data by mutating triplet.

We find that classical D comprises two much stronger but opposing sub-signals (Fig. 5, main text, S5.1, left-hand panel). We note that the exact definition of the major allele has an appreciable impact: D has larger magnitude when calculated over sites where all three non-human hominins carry the same allele. This makes intuitive sense: sites where the non-humans are polymorphic imply variants generated by mutations that occurred outside humans, making them enriched for sites subject to ILS. Similarly, larger magnitude Ds are also seen when the major allele is unambiguous (the same allele inside and outside Africa) compared with using all sites, including those where the African and non-African major alleles differ. This also makes intuitive sense. When the major allele differs between Africans and non-Africans, more sites will be misclassified (in the sense that sites giving positive and negative D will be placed in the wrong category), and some of the signal will be cancelled.

Interestingly, when the same data are plotted after combining the two classes at major-allele sites (i.e., plotting overall or classical D), inter-triplet variation becomes far greater than the variation in either component alone (Fig. S5.1, right-hand panel). The effect is not driven by increased stochasticity, both because sample sizes have increased and because the pattern is highly reproducible across different, independent population pairs (Fig. 2, main text). Consequently, the variation between triplets appears to be an emergent property that reflects a combination of the extent to which the opposing signals cancel each other (ESM8), modulated by the number of D-symmetric sites created when the major allele classes are merged.

To understand these trends, we calculated the overall D for the positive and negative components across a range of simulated conditions (Table S5.1). Without introgression, the negative component, where the Neanderthal allele matches the human minor allele, varies only slightly. In contrast, the positive component, where D is calculated using sites where the Neanderthal allele is the human major allele, increases progressively as the non-African rate drop increases. This is what we expect from the MRVH, where the main driver of D is the class of site generated by recurrent mutations on a BBBBA background. When introgression is added but mutation rate is held constant, the converse is true. Here, the positive component varies little, while the negative component increases with introgression and becomes positive at the highest level.

Direct interpretation of empirical values through comparisons with simulated values is not valid because we do not know either the exact dynamics of the out-of-Africa bottleneck or when any introgression occurred relative to that bottleneck. As we show elsewhere, the split signal is driven by the bottleneck and then modified by whatever drives D. Nonetheless, the two models predict opposite effects on the difference between the positive and negative components: introgression tends to reduce the difference, while mutation-rate variation tends to increase it. Here, the empirical values (positive ~ 0.24, negative ~ -0.14, difference = 0.38) seem more aligned with mutation-rate variation (range 0.311 to 0.448) than with introgression (differences 0.365, 0.265, 0.115 as introgression increases, with 0.265 corresponding to the empirical D) (Table S5.1).

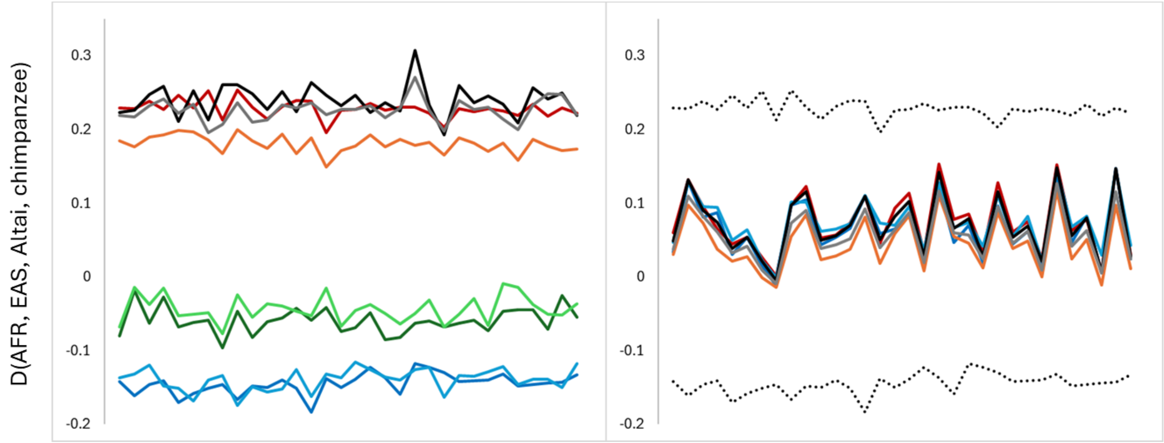

**Figure S5.1. Partitioning D according to whether the Neanderthal carries the human major allele or the human minor allele.** *Data are for transversions only and each represents the average of all possible pairwise population comparisons within each regional comparison*. *The left-hand panel depicts data from several contrasting comparisons: for the major allele = Neanderthal component, lines are EAS-AFR (red), EUR-AFR (grey), SAS-AFR (black), EAS-SAS (orange) and for the major allele = chimpanzee component, lines are EAS-AFR (dark blue), EUR-AFR (green), SAS-AFR (light blue), EAS-SAS (dark green). In all cases, each regional comparison averages all pairwise component comparisons. In the right-hand panel, we combine the two components, revealing the previously described triplet-specific variation in D. Example sub-components are shown as dotted lines for comparison. For the central, combined data plots, we omit the EAS-SAS data, and all remaining lines use the same colors as in the left-hand panel.*

We conclude that partitioning the data by whether the human major allele matches the Neanderthal is not nearly as easy to interpret as we first thought, largely because the out-of-Africa bottleneck splits the signal into positive and negative components. While it would be easy to point to the fact that the negative signal is linked to sites where the Neanderthal allele is the human minor allele and hence goes against the introgression hypothesis, this is far from the whole story because this pattern is only lost if high levels of introgression are simulated. The strongest message is that treating all sites equally can be misleading because it obscures potentially important, opposing signals generated by a combination of the out-of-Africa bottleneck and observation biases, but these opposing signals are not necessarily easy to interpret.

| | POSITIVE | | NEGATIVE | | | | |
|---|---|---|---|---|---|---|---|
| **DROP** | **RISE=1** | **RISE=2** | **RISE=1** | **RISE=2** | **MIG** | **POS** | **NEG** |
| **0** | 0.146 | 0.072 | -0.217 | -0.239 | **0.000** | 0.136 | -0.223 |
| **0.25** | 0.153 | 0.109 | -0.248 | -0.245 | **0.005** | 0.143 | -0.122 |
| **0.5** | 0.176 | 0.138 | -0.261 | -0.233 | **0.010** | 0.140 | -0.047 |
| **0.75** | 0.190 | 0.183 | -0.246 | -0.265 | **0.015** | 0.149 | 0.034 |

**Table S5.1. Partitioning D according to whether the Neanderthal carries the human major allele or the human minor allele: overall D by simulation parameter.** *We simulated a range of scenarios, defined by the drop in mutation rate in non-Africans (0, 0.25, 0.5, 0.75), whether the overall human mutation rate stays constant (RISE=1) or increases twofold linearly over the last ~6,000 generations (RISE=2) and various levels of Neanderthal introgression (0 to 0.015, always with DROP=0 and RISE=1). Positive and negative components correspond to D calculated from sites where the Neanderthal allele matches the human major and minor allele, respectively.*

### ESM6. Variation in D associated with human major allele identity

A key prediction from the standard introgression hypothesis is that a large proportion of introgressed Neanderthal alleles will be rare outside Africa and absent inside Africa (Green et al. 2010). This is because most introgressed material is expected not to be positively selected, and neutral drift acting on rare alleles introduced around 50,000 years ago will rarely have drifted to frequencies above 20% (Green et al. 2010). More alleles could reach high frequency if interbreeding occurred during the out-of-Africa bottleneck, but even then, the proportion would remain modest. Equally, although evidence of introgression within Africa has been reported (Chen et al. 2020), possibly due to low levels of gene flow back into Africa, the amount of Neanderthal material carried by a typical African is thought to be substantially lower than that carried by non-Africans (Bergström et al. 2020).

D depends strongly on the identity of the human major allele (discussed further in ESM8). According to the introgression hypothesis (Green et al. 2010; Durand et al. 2011; Patterson et al. 2012), when the major allele matches the chimpanzee allele (presumed to be the ancestral allele), the remaining minor allele is derived. Sites at which this derived allele is shared with the Neanderthal should be enriched for introgressed variants, generating positive D. By contrast, if the major allele matches the Neanderthal allele, D should be dominated by sites unlikely to have arisen through introgression and tend towards zero. However, we observed the exact opposite pattern (Fig. 6, main manuscript, top-left panel): D becomes large and positive when the major allele matches the Neanderthal allele and large and negative when it matches the chimpanzee allele.

To explore the relationship between D and geography, we compared every 1KG population with every other, calculating D(population1, population2, Altai, chimpanzee) twice: once based only on sites where the major allele matches the Neanderthal and once where it matches the chimpanzee. We summarise the data as boxplots for each major regional comparison (Fig. S6.1). These results mirror the initial finding: when the major allele matches the Neanderthal (top panel), African – non-African comparisons give D in the magnitude range 0.2 – 0.3, with the next largest value being Europe – East Asia; when the major allele matches the chimpanzee, values tend to inverted (previously highest values are now the lowest and *vice versa*) and are generally of lower magnitude. Variability is greatest in comparisons involving Africa and the admixed American populations.

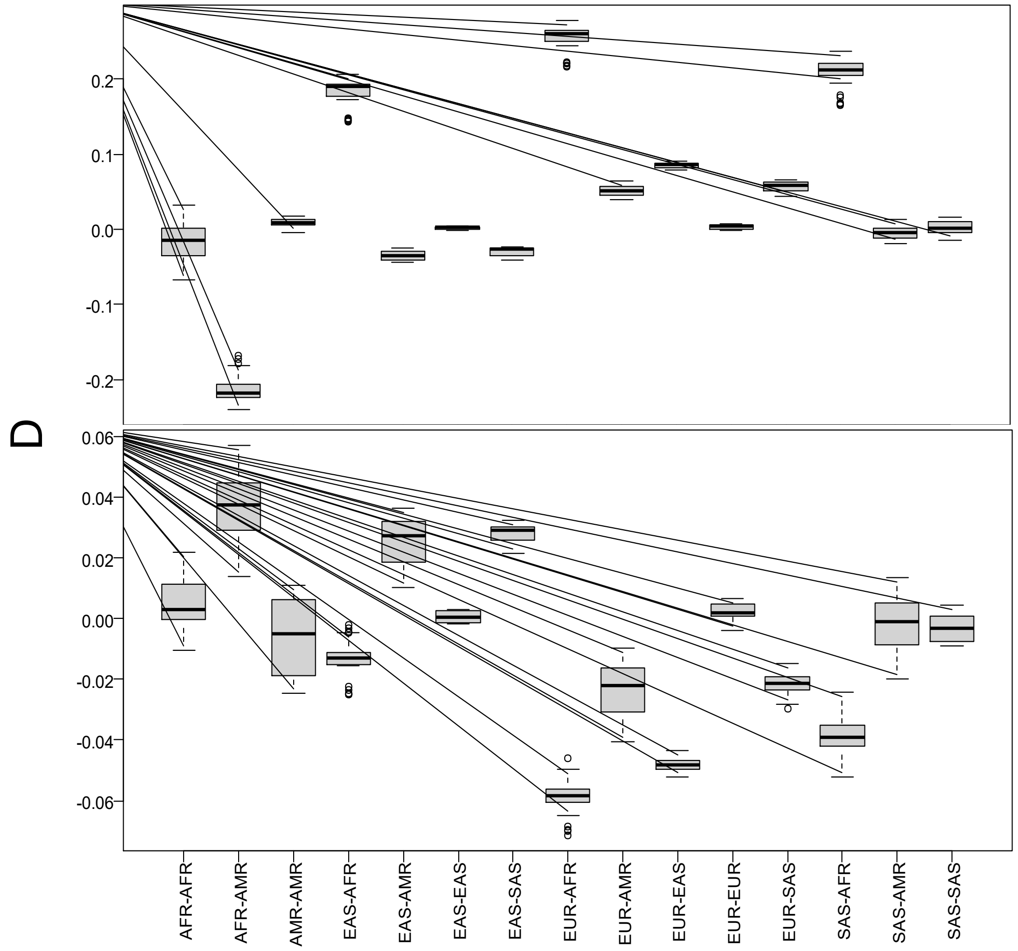

**Figure S6.1. Dependence of D on major allele identity across diverse population pairs.** *Sites contributing to D(population1, population2, Altai, chimpanzee) were categorised into those where the human major allele, defined as having a higher frequency both inside and outside Africa, matched the chimpanzee (top panel) or Neanderthal (bottom panel). Populations were ordered according to how they appear in the downloaded 1KG data files (5 x Europe, 5 x East Asia, 5 x South Asia, 5 x non-admixed Africa, 2 x admixed Africa, 4 x America), and all pairwise comparisons were conducted. D was always calculated the same way, so positive values would classically be interpreted as introgression into population1 (which is why America, appearing after Africa in the list, appears to have an inverted signal).*

Our results are therefore the exact opposite of what is expected under introgression, where a large positive D is expected to be associated with sites where the Neanderthal allele is the minor allele. We note that the definition of the major allele strongly affects the magnitude of D (see ESM8). Here we apply 'unambiguous criteria' by requiring the average frequency in the five non-admixed African populations and in the five European populations both to be above 50%. If, instead of excluding sites where the major allele is unambiguous, we include them by applying less stringent criteria, where the major allele is taken as the allele with the highest frequency across the pooled African and European sample set, the magnitude of D falls by about 30%, although the two sets of D are highly correlated. We assume this is because the closer the frequencies are to 50%, the more sites are misclassified in terms of the 'true' major allele, scrambling site definition and reducing the distinction between classes. In the main text, we applied criteria based on each population pair and obtained a slightly larger D. Again, this presumably reflects how averaging across multiple populations within a geographic region leads to some misclassification, which in turn reduces the magnitude of D.

More broadly, our results suggest that D is not simply measuring the abundance of introgressed Neanderthal alleles, as is commonly assumed. Instead, D is strongly influenced by the underlying properties of D-informative sites, including their frequency spectra and the processes that shape them. That the signal runs opposite to what the standard introgression model predicts implies that D captures a broader feature of population genetic variation than introgression alone. Consequently, D must not be interpreted uncritically as an unambiguous measure of Neanderthal ancestry.

Understanding what D measures requires identification of the mechanisms that generate its signal. We argue that these mechanisms generate patterns that are inconsistent with the basic expectations of the introgression hypothesis. We show that D is strongly associated with heterozygosity differences between populations and with the identity of the major allele, both of which profoundly influence its magnitude and direction (e.g., Figs. S4.5, S5.1, S6.1). These dependencies persist across alternative definitions of the major allele and remain evident even when analyses are restricted to site classes that, under the standard introgression model, should contribute little or no signal. D appears to be a composite statistic that reflects broader features of population genetic variation. Any interpretation of D as evidence for introgression must therefore first account for these dependencies and demonstrate that the observed signal cannot be generated by the demographic and mutational processes that also shape heterozygosity and allele frequencies.

### ESM7. Realistic forward simulations.

The SLiM package (Haller, Ralph, and Messer 2026) is extremely flexible and allows realistic simulation of complex demographic and mutational models. We based our simulations on the default demography model generated by Gravel et al. (2011), available as a 'recipe' in SLiM's manual, adding in chimpanzee (*N*=50) and Neanderthal (*N*=5000) lineages (methods). Population sizes were chosen to maximise efficiency: the small chimpanzee population in particular reflects a need to minimise the number of polymorphic sites being tracked when we are only interested in whether a mutation has or has not occurred on the lineage leading to a single sampled chimpanzee; larger numbers of segregating variants in the chimpanzee greatly increase run times but add no extra realism to the output. Mutation rates were simulated either as a simple Kimura model ("ONEMU"), with transitions four times as likely as any given transversion, or as a full mutation spectrum model with each of the 192 possible mutations given its own rate ("TRIPLETS", 64 triplets x 3 possible substitutions), estimated from empirical data. In both models, we normalized mutation rates to an average of $2.36 \times 10^{-8}$ (Gravel model). Because the Gravel demography generates a realistic out-of-Africa bottleneck, we cannot use it directly to simulate weaker bottlenecks or no bottleneck. For these conditions, we replaced the non-African size-through-time profile with a size of 10,000 that is either constant (= no bottleneck) or reduced to some smaller size for 50 generations around 2,000 generations before the present (= "step bottleneck"). Introgression was modelled by adding a pulse of migration immediately after the step bottleneck. Note that when introgression is added to the Gravel model, where the bottleneck is long and slow, introgressed material enters humans near the start of the bottleneck, allowing it to be subject to much greater genetic drift. This distinction has important consequences (see below) that have not previously be discussed. We applied mutation-rate changes across all rates and tested three levels of rate reduction in non-Africans relative to Africans (0.25, 0.5, and 0.75), with or without a general linear increase in mutation rate in humans over the last ~6,000 generations.

#### ESM7.1 Simulated D across the mutation spectrum

We first tested a null model with no introgression and no variation in mutation rate across humans. We were surprised to find that, although D never deviates significantly from zero, there is a weak patterning across the mutation spectrum (Fig. S7.1). Here, we ran 1000 simulations, each modelling a 500Kb segment, in four sets of 250 replicates (different-coloured lines). While there is much stochasticity, there is clear similarity between the four tracks, particularly for transitions and for the most mutable XCG triplets (the four largest peaks in ONEMU_TS). In both cases (ONEMU and TRIPLETS), D calculated over all 1000 simulations is essentially zero (-0.001 +/- 0.01 s.e.m., -0.003 +/- 0.007 s.e.m., respectively).

The similarity between replicates is surprising because the theoretical framework developed by Durand et al. (2011) indicates that as long as the mutation rate is constant across populations, D should not vary with mutation rate. Our results suggest either a minor bug in our SLiM implementation or an incorrect assumption in the theory. To test which explanation is more likely, we calculated correlation coefficients between the triplet-specific D values in each of the four replicate blocks for the TRIPLETS scenario and the equivalent empirical D values, obtaining $r^2$ values of 0.38, 0.37, 0.19, and 0.11 (all $p<0.01$). Thus, while the overall simulated D is zero, individual triplets give D values that covary with the D values in the real data spectrum (Fig. S7.2), suggesting that at least part of the variation in D across triplets is an emergent property that reflects the only property directly shared between the simulations and real data, namely the way mutation rates vary between triplets. We do not yet understand the underlying mechanism, though it may reflect non-linearities at mutation hot spots where the most mutable triplets locally deplete the number of unmutated sites. This warrants further investigation.

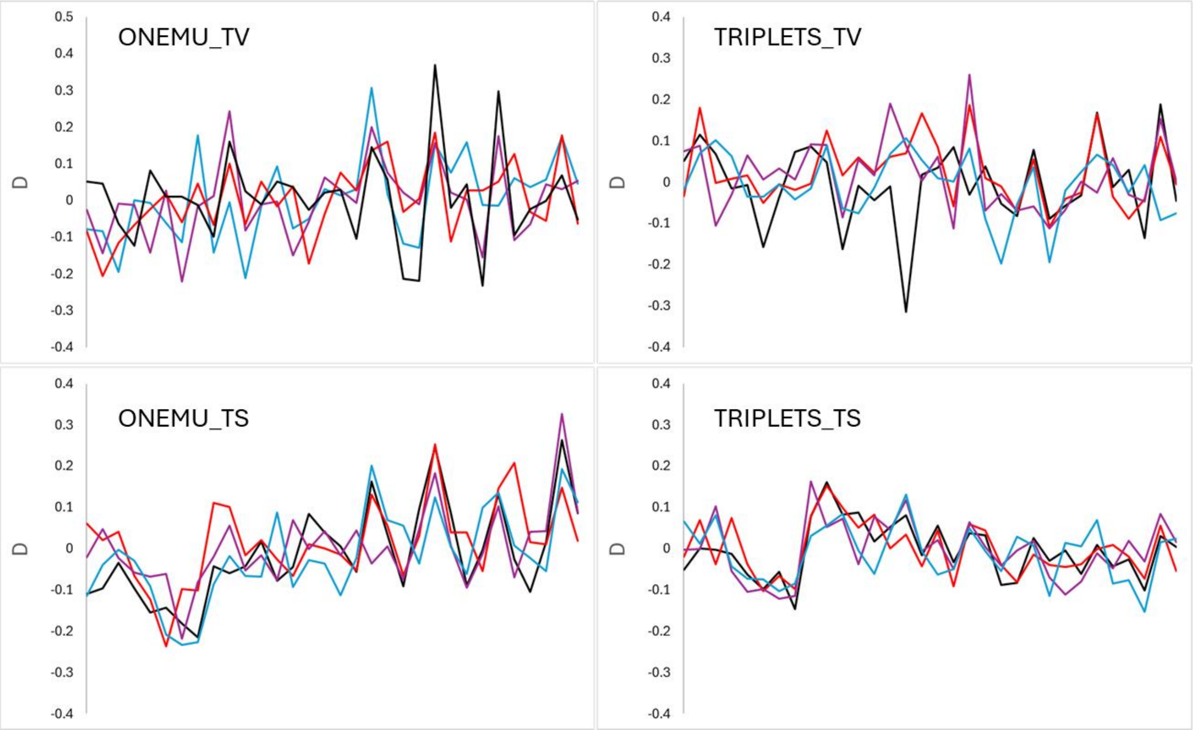

**Figure S7.1. Variation in D in simulated data where mutation rates do not vary across populations.** *Each line represents pooled data from 250 independent runs, each modelling the fate of a randomly selected 500Kb segment from chromosome 1. Two mutation models were used: ONEMU uses one mutation rate for transitions and one for transversions; TRIPLETS uses separate mutation rates for all 64 triplets × 3 possible substitutions.*

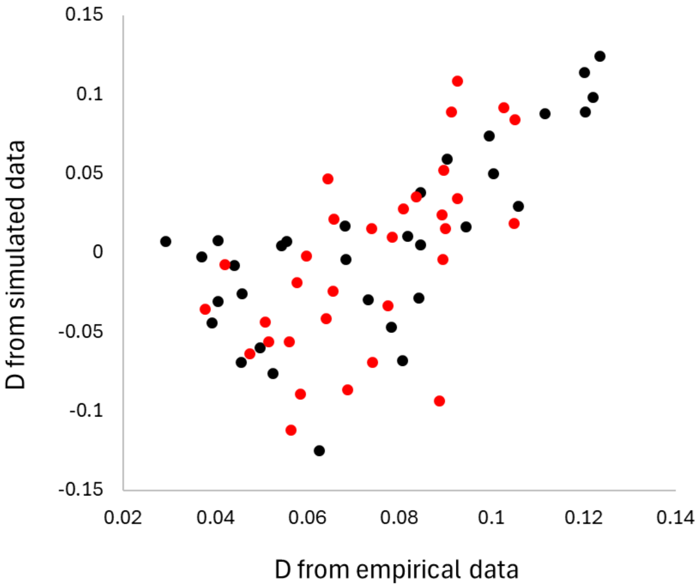

**Figure S7.2. Correlation between empirical and simulated triplet-specific D.** *Both transversion (black) and transitions (red) exhibit similar positive correlations, despite the simulations not including either introgression or variation in mutation rate.*

### ESM7.2 Triplet-specific variation in D under the two competing hypotheses

To explore fit to the MRVH, we simulated a 75% drop in mutation rate among non-Africans. Replicating the constant mutation-rate simulations, triplet-specific D values are significantly positively correlated (Student *t*-test, $N$=64, $r^2$=0.43, $p$<<0.0001) with those from empirical data, but the relationship is even stronger (Fig S7.3, left-hand panel). We also asked whether it related to mutability, finding a significant correlation with the expected number of mutations at each triplet, estimated as mutation rate times the number of sites in the genome. Puzzlingly, the correlation is negative (Fig S7.3, right-hand panel). While we do not yet fully understand this, it should be remembered that D reflects a subtle balance between the number of sites being generated that are D-asymmetric, thereby tending to increase D, and those that are D-symmetric, acting to decrease D. Higher mutation rates tend to increase both classes, so while it may seem 'obvious' that higher mutation rates will drive larger D values, this is not necessarily so. Indeed, the highest and lowest triplet-specific D values tend to occur for particular transversions, not among transitions, which occur with higher probability (Fig. 2, main text). Notably, XCG transversions give very high D, yet the most mutable sites overall, XCG transitions, give rather average values.

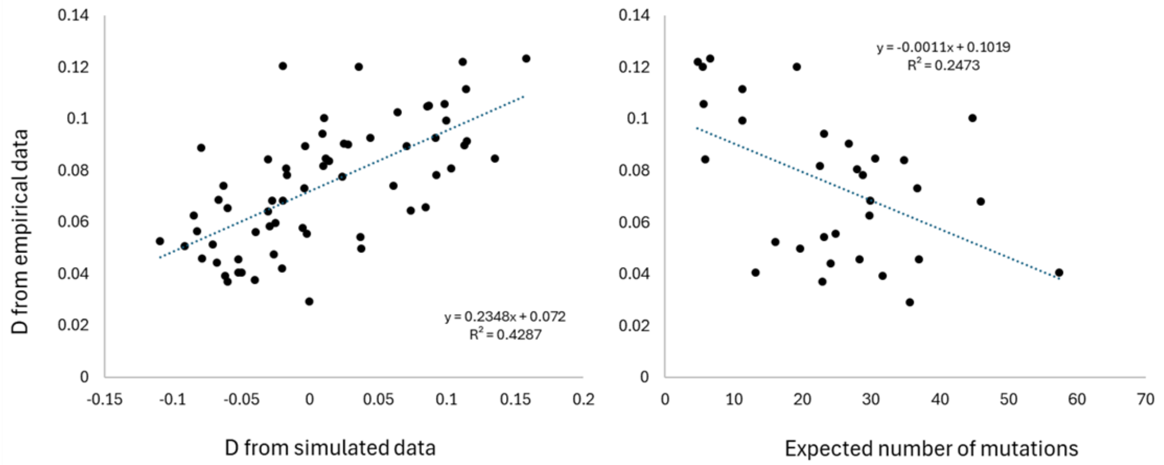

**Figure S7.3. Relationship between empirical triplet-specific D, simulated data and the expected number of mutations.** *The left-hand panel plots triplet-specific D from simulated data (x-axis) against equivalent values from real data, revealing a significant positive correlation (N=64, $r^2$=0.43, p<<0.0001). The right-hand panel depicts the relationship between the expected number of mutations at each triplet (mutation rate × genome-wide frequency) and empirical triplet-specific D, showing a significant negative trend for transversions (N=32, $r^2$=0.25, p=0.004), where variation in D is greatest. However, the trend becomes marginally non-significant overall (N=64, $r^2$=0.053. p=0.067).*

To test fit to the introgression hypothesis, we simulated 1.5% introgression, giving an overall D of 0.11. We chose this large value so even weak trends would be revealed. As before, D does appear to vary between triplets. However, this variation is entirely uncorrelated with the triplet-specific D values from empirical data (Fig. S7.4). We note that the XCG triplets form an outlying group of four points at the bottom of the graph. This is consistent with other analyses in which XCG triplets yield either very large values, when their high mutation rates dominate, or very low values when, as here, their general rarity in the genome more than compensates for the potentially large signal each might give per site. That the simulated introgression values are uncorrelated with empirical values argues strongly that introgression is not the dominant force driving D. Instead, our analyses suggest that the way mutation rates and local frequencies vary across triplets tends to generate predictable variation in D that is then effectively erased when D is dominanted by introgression: introgressed fragments carry triplets in proportion to their genomic frequencies which effectiewly scrambles the underlying pattern that has a more complicated origin.

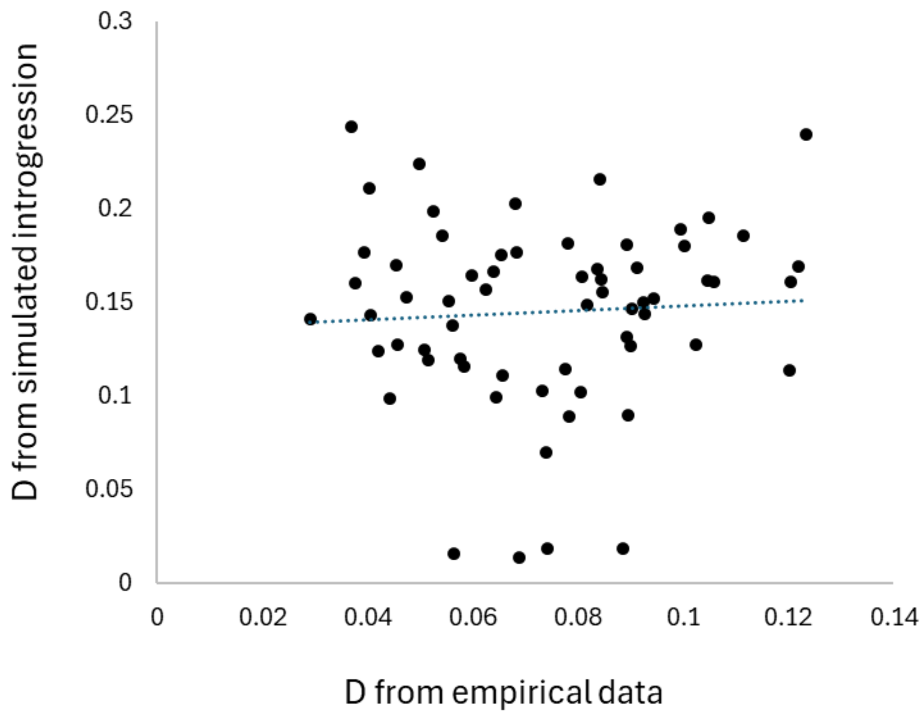

**Figure S7.4. Triplet-specific D from simulated introgression is uncorrelated with empirical triplet-specific D.**

**ESM7.3 D and relative mutation rate under different simulation model parameters.**

To explore the behaviour of D and our measure of relative mutation rate, we ran a series of simulations. We focused mainly on the Gravel model and ran a default number of 500 replicates for each set of parameter values, each simulating the fate of a randomly selected 500Kb segment drawn from the human reference sequence for Chromosome 1. We simulated the following conditions:

a) Mutation model; either a full triplet model where all triplets have their own mutation rates, estimated from empirical data (TRIPLETS) or a simple Kimura model with a different rate for transversions and transitions (ONEMU).
b) Non-African rate reduction. Mutation rates in non-Africans were reduced at their formation to 75%, 50% or 25% of the African value.
c) Mutation rate rise. We speculated that human mutation rates might have been generally rising. We simulated two states: no rise (RISE=1) and a linear rise starting from 6000 generations ago to a final rate of double that in other lineages (RISE=2) .

Our conditions are far from exhaustive but cover enough parameter values to infer which are likely compatible with empirical values. Results are summarised in Table S7.1.

| | | Triplets | | OneMU | |
|---|---|---|---|---|---|
| **Drop** | **Rise** | **MU** | **D** | **MU** | **D** |
| **0** | **1** | 0.998 | 0.01 | 1.000 | -0.001 |
| **0** | **2** | 0.990 | -0.003 | 0.995 | -0.003 |
| **0.25** | **1** | 0.960 | 0.002 | 0.964 | -0.0035 |
| **0.25** | **2** | 0.922 | 0.023 | 0.927 | 0.010 |
| **0.5** | **1** | 0.923 | 0.011 | 0.930 | 0.016 |
| **0.5** | **2** | 0.850 | 0.046 | 0.850 | 0.030 |
| **0.75** | **1** | 0.881 | 0.022 | 0.884 | -0.004 |
| **0.75** | **2** | 0.779 | 0.07 | 0.777 | 0.033 |

Table S7.1. **Variation in simulated D and relative mutation rate across a range of parameter values.** *Model conditions are described above.*

Several important trends are apparent. First, when Drop=0, D is indistinguishable from zero and relative mutation rate is indistinguishable from unity. Second, as Drop increases from a 25% rate reduction (Drop=0.25) to a 75% rate reduction (Drop=0.75), relative mutation rates drop progressively below one while D increases. Third, increasing the human mutation rate has a large impact, exaggerating the effect on both D and the relative mutation rate. This makes sense because a rising mutation rate pushes a greater proportion of derived variants towards the present, thereby increasing the impact of a recent drop in the non-African rate. Finally, and arguably the most important trend, while the TRIPLET and ONEMU models give essentially identical values for relative mutation rate (as they should), D is consistently much larger in the TRIPLET model. This observation emphasises the importance of recurrent mutations. In the full TRIPLET model, the range of mutation rates across sites is much higher than in the ONEMU model, despite the same mean. Since the rate of recurrent mutations scales with the mutation rate squared, recurrent mutations are far more likely in the full TRIPLET model. This confirms our expectation that D generated under the MRVH increases with variation in mutation rates across sites through recurrent mutations. Moreover, we simulate only inter-triplet variation in rate, but in nature there is also massive variation in mutation rate along each chromosome, which will further enhance these effects. Even without the contribution of mutation hot spots, realistic values of D of ~0.05 can be generated by a modest 50% reduction in mutation rate, provided that the human mutation rate has increased over time.Factoring in mutation hot spots will generate a larger effect size, either removing the need to invoke a rising rate in humans, or reducing the size of the non-African drop in rate.

### ESM8. Challenges in defining the human major allele

Partitioning data according to which allele is the major or minor allele offers a critical tool for unpicking the mechanism(s) that underpin D. In general, the major allele will tend to be the ancestral state while the minor allele will tend to be the derived state, reflecting recent processes, such as introgression or changes in mutation rate. Unfortunately, defining the human major allele is far more complicated and prone to bias than it may at first appear (e.g., Keightley and Jackson 2018). We have explored several options and have had to accept that none are perfect or bias-free. Moreover, the choice of method impacts the downstream results. In the following section, we outline the problems and how we have tried to mitigate the worst.

A large majority of minor alleles are very rare and will be classified as minor under any set of criteria. However, very rare alleles tend to contribute little to most metrics, including D, because they are, by definition, absent from almost all haplosomes. In contrast, sites with two alleles at roughly equal frequency are rare, but a high proportion of haplosomes carry whichever allele is critical in each analysis (major/minor). Moreover, when the two alleles are present at around equal frequency, the assignment of major/minor labels becomes effectively arbitrary, because genetic drift, sampling biases, and other factors exert far more influence than the identity of the ancestral state. One tactic might be to exclude sites where the identity of the major allele is ambiguous. For example, we might require the major allele to be the same both inside and outside Africa. However, this approach does not work for mutation rate estimation, where the method is only valid and robust to differences in demographic history if all alleles are considered. Methods that require allele filtering also risk introducing bias because African and non-African allele frequency spectra differ.

Similarly, the reliability of the assumption that the major allele is the ancestral state also varies with minor allele frequency. Extremely rare alleles not introduced by admixture almost certainly represent derived states. However, as minor allele frequency increases, the chance that the minor allele is the ancestral state rises to around 50%, when the classification is arbitrary. It follows that biases due to sample size and sample structure, which dictate the relationship between 'true', underlying allele frequency and the frequency in the sample of genomes being studied, will interact with these uncertainties to confound the estimation of the ancestral state further. The out-of-Africa bottleneck further complicates the situation, where drift has a much greater impact (Henn et al. 2015; Ashraf and Lawson 2021). Indeed, the bottleneck's extreme impact is illustrated by the fact that in simulations without a bottleneck, D no longer splits into positive and negative components (ESM5 above).

A second issue relates to the nature of rare alleles. As stated, individual very rare alleles contribute negligibly to most analyses (e.g., D). However, as a large class, they dominate analyses based on counts of segregating sites, such as mutation-spectrum analyses. This impact is especially large when counting how many times each triplet has mutated, where a site carrying a rare allele counts the same as one with a common allele. Consequently, such analyses are highly sensitive to sample size and sampling strategy. Large sample sizes increase the probability of detecting rare variants, while samples drawn from a narrow geographic range have the opposite effect, because any level of relatedness between sampled individuals will act to reduce the number of truly independent alleles sampled.
